# Mapping of the fibrinogen-binding site on the staphylocoagulase C-terminal repeat region

**DOI:** 10.1101/2021.06.29.450373

**Authors:** Ashoka A. Maddur, Markus Voehler, Peter Panizzi, Jens Meiler, Paul E. Bock, Ingrid M. Verhamme

## Abstract

The N-terminus of *S. aureus* staphylocoagulase (SC) triggers activation of host prothrombin (ProT), and the SC·ProT* complex cleaves host fibrinogen (Fbg) to form fibrin (Fbn) deposits, a hallmark of SC-positive endocarditis. The C-terminal domain of the prototypical Newman D2 Tager 104 SC contains 1 pseudo-repeat (PR) and 7 repeats (R1→R7) that bind Fbg/Fbn Fragment D (Frag D). This work defines affinities and stoichiometries of Frag D binding to single- and multi-repeat C-terminal constructs, using fluorescence equilibrium binding, NMR titration, Ala scanning, and native PAGE. Constructs containing PR and each single repeat bound Frag D with *K*_D_ ~50 - 130 nM and a 1:1 stoichiometry, indicating a conserved binding site shared between PR and each repeat. NMR titration of PR-R7 with Frag D revealed that residues 22-49, bridging PR and R7, constituted the minimal peptide (MP) required for binding, corroborated by Ala scanning, and binding of labeled MP to Frag D. MP alignment with the PR-repeat and inter-repeat junctions identified conserved residues critical for binding. Labeled PR-(R1→R7) bound Frag D with *K*_D_ ~7 - 32 nM and stoichiometry of 1:5; and PR-R1R2R3, PR-R1R6R7, PR-R3R4R7, and PR-R3R6R7 competed with PR-(R1→R7) for Frag D binding, with a 1:3 stoichiometry and *K*_D_ ~7 - 42 nM. These findings are consistent with binding at the PR-R junctions with modest inter-repeat sequence variability. Circular dichroism of PR-R7 and PR-(R1→R7) suggested a largely disordered structure and conformational flexibility, allowing binding of multiple fibrin(ogen) molecules. This property facilitates pathogen localization on host fibrin networks.

## Introduction

*Staphylococcus aureus* (*S. aureus*) is the leading cause of hospital-acquired infections targeting the skin, soft tissue, bone, and heart valves. *S. aureus* can cause endocarditis, osteomyelitis, and septicemia (1). Invasive methicillin-resistant infections are responsible for ~20,000 deaths annually (2). The *S. aureus* staphylocoagulases (SC) are secreted virulence factors, originally grouped into 12 serotypes based on their genetic diversity (3), and with recent serotype additions now a total of 15 (4,5). SC contributes to abscess formation, typical of *S. aureus* infection (6–8). Interaction of host fibrinogen (Fbg) with the *S. aureus* virulence factors, clumping factor A and B, FbnA, efb and SC was reported previously (9–13), however several mechanisms underlying these specific Fbg interactions are not yet defined at the molecular level. Coagulase-positive *S. aureus* adheres to exposed subendothelium on heart valves damaged by turbulent blood flow, where deposition of platelets and fibrin (Fbn) creates adhesion foci for circulating bacteria (1). Fbg also facilitates interactions between *S. aureus* and platelets (14). Our previous fluorescence equilibrium binding and native PAGE studies revealed for the first time that Fbg and Fbn bind free SC as well as its complex with host prothrombin (ProT); and that the Fragment D (Frag D) domain of Fbg/Fbn harbors major interaction sites for binding to the C-terminal repeat region of SC (15,16). The current work identifies the binding motifs in the C-terminal SC region, essential for interaction with Frag D, and defines the stoichiometry of Frag D binding to free and ProT-complexed SC.

SC is a bifunctional protein that uses its N-terminus for conformational activation of prothrombin (ProT), the precursor of the central clotting protease, thrombin (T). Classical serine proteinase activation requires proteolytic cleavage of the Arg^15^-Ile^16^ peptide bond in the zymogen (chymotrypsinogen numbering), with release of a new N-terminus that inserts in the Ile^16^ binding pocket to trigger formation of the active site (17,18). In contrast, our structure-function studies showed that formation of the active site in ProT by SC is a non-proteolytic mechanism, in which SC inserts its N-terminus into the zymogen activation pocket, thereby forming a critical salt bridge required for conformational expression of the active site in ProT (19,20). In the absence of SC, Fbg binds to thrombin at exosite I and is cleaved as a substrate. In the SC·ProT* (* denotes a fully expressed active site) and SC·thrombin complexes, (pro)exosite I is occupied by the tight-binding N-terminal D2 domain, and a new binding site is expressed on these complexes for Fbg as a substrate (21). The SC(1-325) fragment, containing the D1 and D2 domains, is sufficient for expression of this novel Fbg binding site on the SC·ProT* and SC·thrombin complexes, and triggering of proteolytic Fbg cleavage to form Fbn clots.

In addition to the N-terminal D1 and D2 ProT-binding domains, the prototypical full-length SC(1-660) from the *S. aureus* strain Newman D2 Tager 104 also contains a central region with unknown function, and a C-terminal domain consisting of one 32-residue pseudo-repeat (PR) and seven 27-residue tandem repeats, R1→R7 (**Fig. 1**) with highly conserved inter-repeat sequences. SC from other strains contains the PR and 4 to 9 repeats (**Fig. S1**) (22,23). Fbg binding to SC was recognized previously (14,24,25), and we narrowed down this interaction to Fbg Fragment D (Frag D) binding to the C-terminal domain (15,16,26,27). Binding of Fbg and Frag D to the SC C-terminal domain has been studied by turbidity assays, ELISA, isothermal titration calorimetry and solid-phase fibrinogen binding of an SC C-terminal domain construct (25,28,29). The present work identifies a minimal C-terminal repeat region, or minimal peptide (MP), required for binding of PR-R constructs to Frag D, by equilibrium binding, NMR titration, and alanine scanning. The MP sequence bridges the PR and R junctions, exhibits alignment with inter-repeat junctions, and contains a set of conserved residues that are critical for Frag D binding. Knowledge of this minimal binding site is useful for future development of mechanism-based antibody drugs targeting the SC C-terminal domain.

**Figure 1:**
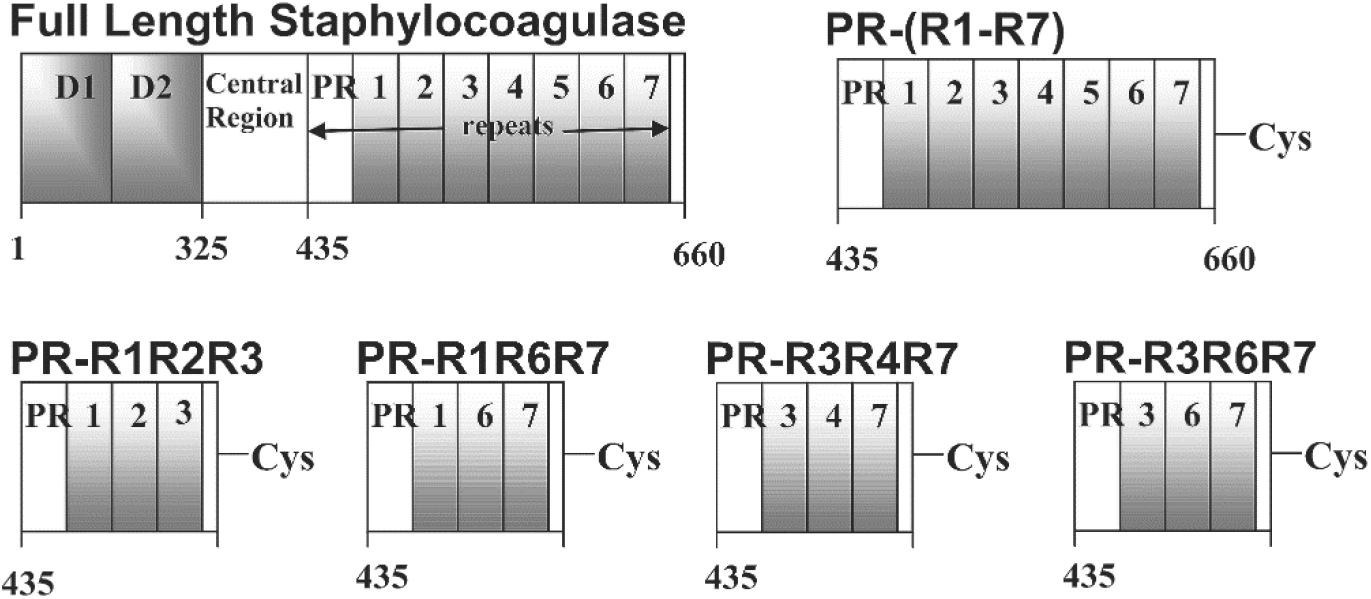
Domain organization of full-length SC, and multi-repeat constructs. Full-length SC secreted by the *S. aureus* Newman D2 Tager 104 contains 3 major regions: the N-terminal D1 and D2 domains (I), a central region (II), and the C-terminal repeat region (III), consisting of 1 pseudorepeat, PR, and 7 repeats R1-R7. The 5 residues at the C-terminal end are conserved (IV).

We also present an in-depth analysis of the potential of SC to bind multiple Frag D units at its C-terminal repeat domain, independent of substrate Fbg binding to the catalytic SC·ProT* complex. We show that the full-length C-terminal repeat construct, PR-(R1→R7) binds Frag D with a stoichiometry of ~5 Frag D per construct; and constructs containing a leading PR and three R units in consecutive or arbitrary order, bind Frag D with a stoichiometry of ~3. Shortening of the full-length repeat construct or changing the order of repeats did not alter the ability of repeats to bind Frag D, reaffirming the requirement of aligned conserved residues at the repeat junctions for formation of multiple binding sites. Consistent with these observations, our data of *in vivo* localization of SC·ProT* complexes to areas rich in Fbg involve a template mechanism that utilizes the SC C-terminal domain (15). The high affinity of multiple binding sites for Fbg/Fbn D domains on a single SC molecule contributes to a mechanism of selective anchoring of proteolytically active SC·ProT* complexes on host fibrin-bacteria vegetations subjected to arterial shear stress.

## Results

### Sequence alignment of the SC C-terminal repeats

The C-terminal repeat alignment of SC(1-660) *S. aureus* Newman D2 Tager 104 was performed using an online box shade alignment program, http://embnet.vital-it.ch/software/BOX_form.html. **Figure 2** shows the sequence alignment of PR, 32 amino acid residues, with R1 to R7, each 27 residues in length. PR and the repeats show a 30-40 % conserved sequence, and there is >80 % pairwise identity among the repeats. Repeats R2, R4 and R5 are identical. We identified the MP sequence as described below, and in **Figure 13** the MP alignment is shown with the bridging sequences between repeats, and conserved residues (*red*) at positions G^37^, Y^41^, A^43^, R^44^, P^45^, K^49^ and P^50^ (PR-R7 numbering). The residues G^28^ and I^31^ are present only in the PR, and are replaced with similar size hydrophobic residues A and V in the repeats.

**Figure 2:**
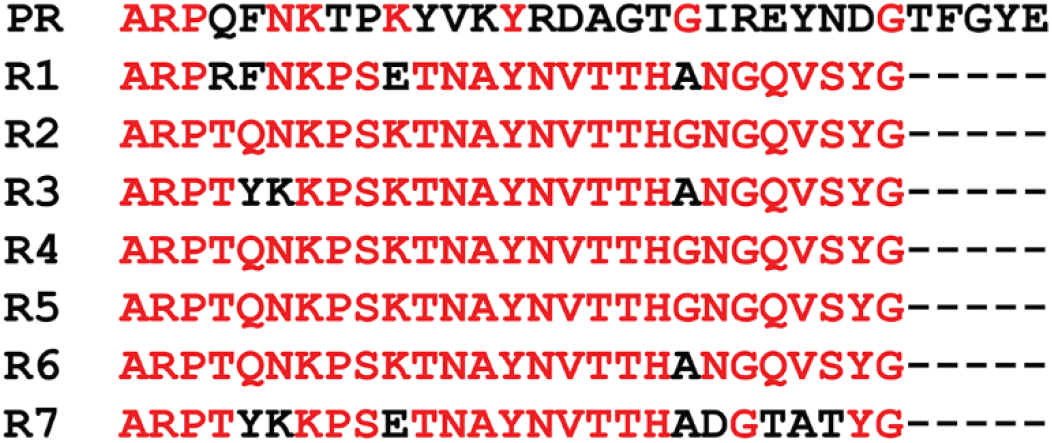
Sequence alignment of the C-terminal pseudo-repeat PR and the repeats R1 - R7 from SC of *S. aureus* Newman D2 Tager 104. The PR contains 32 residues, and the repeats R1-R7 are 27 residues in length. Conserved residues are in red.

### Size exclusion chromatography and light scattering

SC(1-325) containing only the N-terminal domains D1 and D2, and full length SC(1-660) that includes the C-terminal repeats, were chromatographed separately and in mixtures with Frag D. **Figure 3A** shows the elution profile of SC(1-325) (*black*), Frag D (*red*), and their mixture (*blue*). Light scattering indicated that SC(1-325) (M_r_ 38,000 Da) eluted with an apparent molecular mass of ~32,500 Da. Frag D (M_r_ 88,000 Da) showed a hydrodynamic property-related anomaly, eluting in the same apparent molecular mass range, in part due to interaction of Frag D with the column matrix. The SC(1-325) and Frag D mixture eluted at a position similar to SC(1-325) and Frag D alone, suggesting that SC(1-325) does not form a high M_r_ complex with Frag D. **Fig. 3B** shows the elution profile of SC(1-660) (*black*), Frag D (*red*), and their mixture (*blue*). Light scattering indicated that SC(1-660) (M_r_ 74,390 Da) eluted with apparent molecular mass of ~81,200 Da. The mixture of SC(1-660) and Frag D showed a new peak eluting at ~165,000 Da, suggesting that the C-terminal region of SC(1-660) binds Frag D.

**Figure 3:**
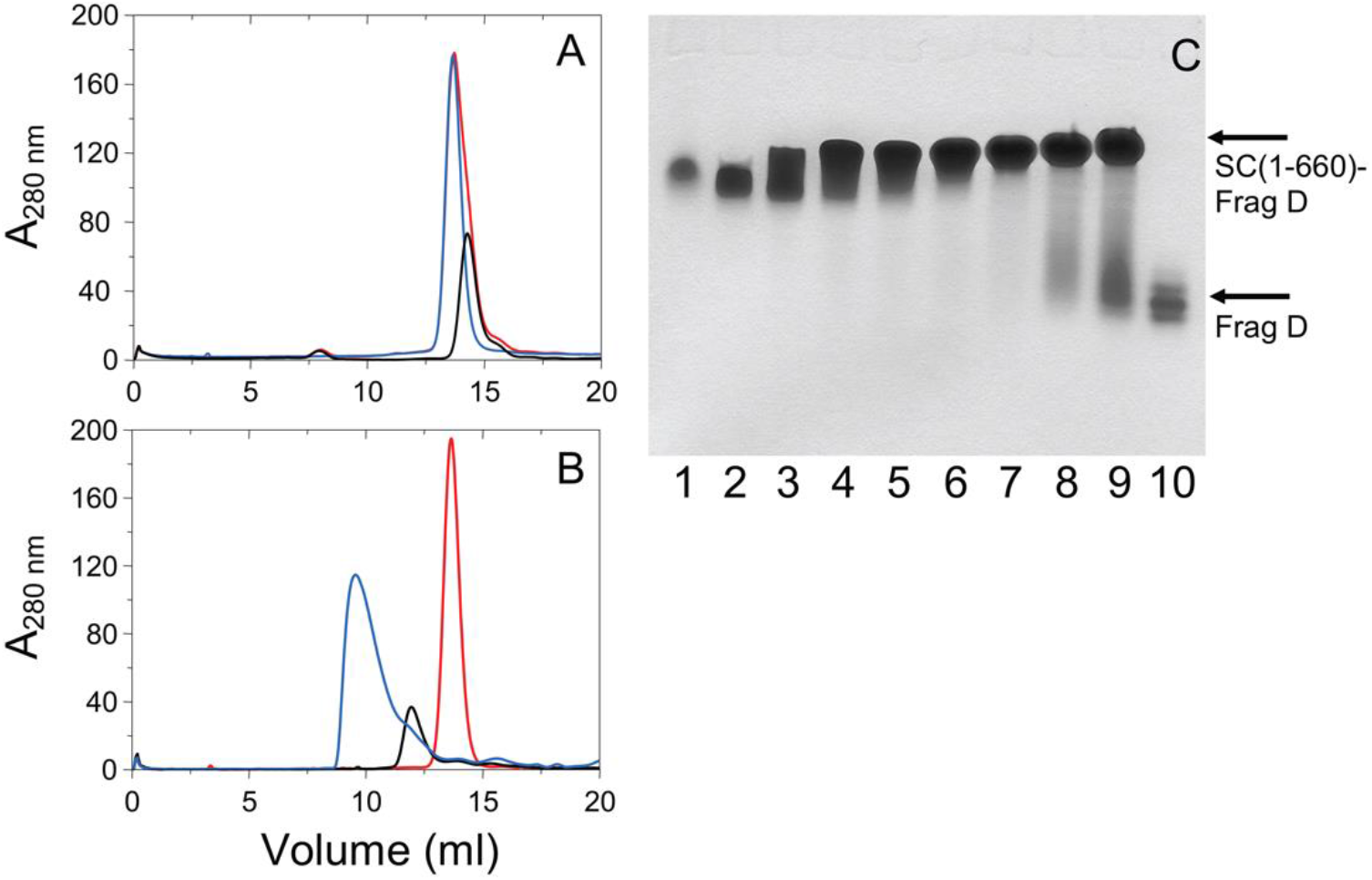
Size exclusion chromatography of SC(1-325), SC(1-660) and their mixtures with Frag D, and native PAGE of SC(1-660) binding to Frag D. *A*, elution profiles (A_280 nm_ in mOD) of SC(1-325) alone, *black*; Frag D alone, *red*; mixture of SC(1-325) and Frag D, *blue*. *B*, elution profiles of SC(1-660) alone, *black*; Frag D alone, *red*; mixture of SC(1-660) and Frag D, *blue*. SC(1-660) forms a high molecular weight complex with Frag D as shown by the appearance of new high molecular weight peak. *C*, Coomassie-stained native 6 % Tris-Glycine PAGE, showing the formation of SC(1-660)·Frag D complex. SC(1-660) (4.0 μM) was reacted with Frag D (lanes 2-9 at respectively 4.1, 8.2, 16.4, 20.0, 24.6, 28.6, 32.7 and 36.0 μM) at 25 °C for 30 minutes before separation at 4 °C. Lanes 1 and 10 are SC(1-660) and Frag D controls, respectively.

### Native PAGE of Frag D binding to SC(1-660), [5F]PR-R1, the SC(1-660)·[5F]ProT complex, and labeled C-terminal repeat constructs

Formation of multimeric SC(1-660)·Frag D complexes was confirmed by native PAGE (**Fig. 3C**). Frag D (lane 10, M_r_ 88 KDa) runs faster than SC(1-660) (lane 1, M_r_ 74.4 KDa) due to combined differences in isoelectric point, hydrodynamic properties, secondary structure, and native charge density, unlike in SDS-PAGE. Minor Frag D proteolysis bands were due to fibrinogen digestion by trypsin during Frag D preparation. Upon binding of Frag D to SC(1-660), the SC(1-660)·Frag D complex band shifted, starting at 2-fold molar excess Frag D, and the shift and band intensity became more pronounced at increasing Frag D (lanes 2-9). Unbound Frag D was observed starting at 7-fold molar excess (lanes 7-9), suggesting that the C-terminal region of SC(1-660) binds multiple molecules of Frag D, possibly 5 to 6.

Frag D also formed a complex with PR-R1, labeled with 5-iodoacetamidofluorescein (5-IAF) at an engineered C-terminal Cys residue ([5F]PR-R1), but not with labeled R1 peptide ([5F]R1) or pseudo-repeat ([5F]PR) (**Fig. S2**).

Tight-binding complexes of SC(1-660) with ProT, labeled at the active site with 5-fluorescein ([5F]ProT), recruited multiple Frag D molecules at the C-terminal SC domain. In incubations of 2.5 μM [5F]ProT with 3.75 μM SC, [5F]ProT binding was saturated due to the tight *K*_D_ of this interaction (30). Subsequent incubation of the complex with molar excess Frag D yielded higher order complexes (**Fig. 4AB**). All the [5F]ProT was bound in the SC(1-660)·[5F]ProT binary complex, with 1:1 molar stoichiometry (lane 3). Binding to multiple Frag D molecules occurred, with unbound Frag D observed at 8-fold molar excess (lane 9).

**Figure 4:**
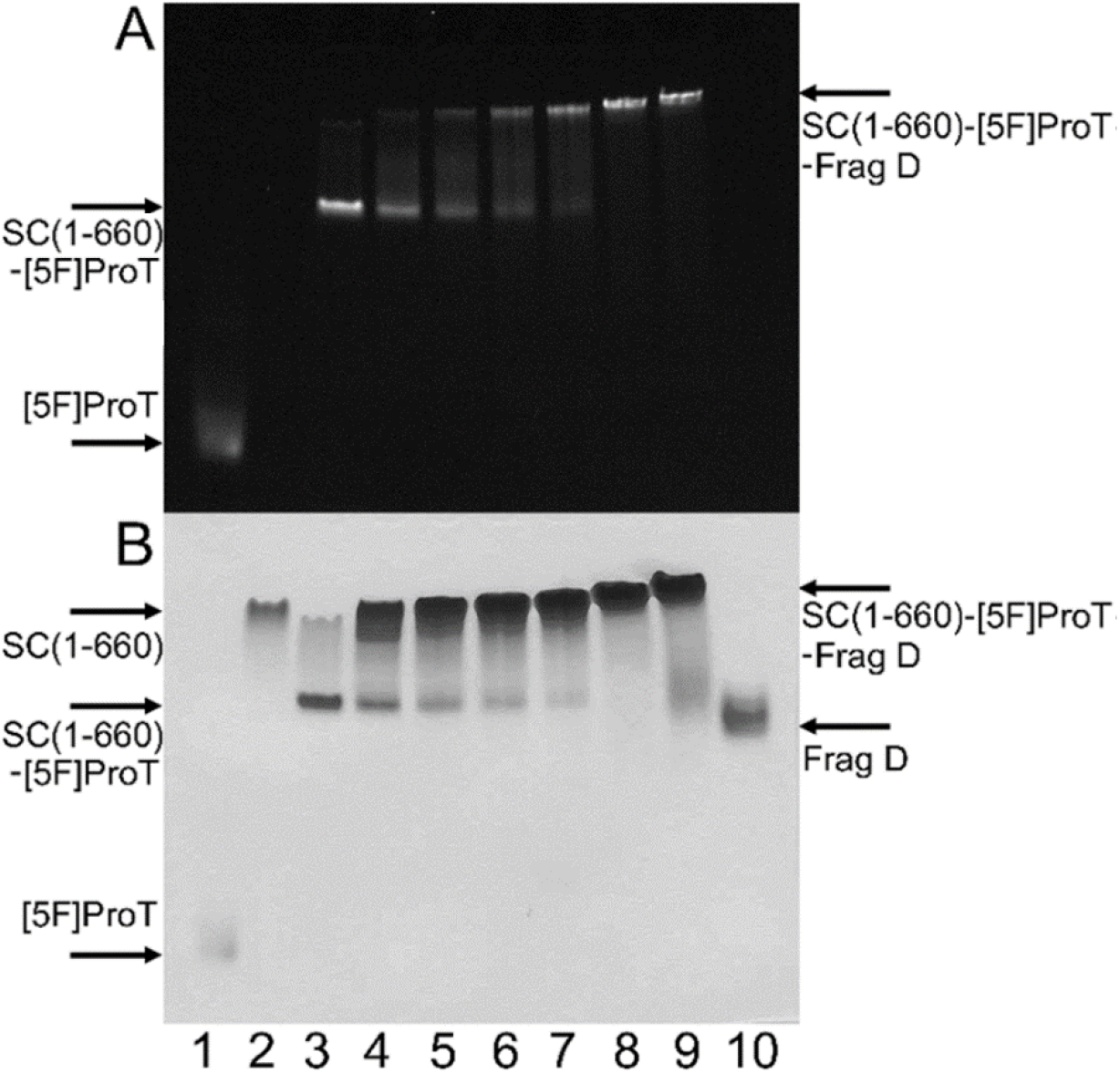
Binding of Frag D to the SC(1-660)·[5F]ProT complex: ***A***, fluorescence and ****B****, Coomassie stain. Incubation of [5F]ProT (2.5 μM) with SC(1-660) (3.7 μM) for 30 minutes, and formation of the SC(1-660)·[5F]ProT binary complex (lane 3). The complex was reacted with Frag D (lanes 4-9, respectively 7.5, 11.2, 15.0, 18.7, 26.2 and 29.9 μM) at 25 °C for 30 minutes to form higher order complexes. Lanes 1, 2 and 10 are [5F]ProT, SC(1-660), and Frag D controls. Proteins were run on a 6 % Tris-Glycine native gel at 4 °C.

Frag D bound to [5F]PR-(R1→R7) containing all seven repeats in order (**Fig. 5AB**). Higher order complex formation was indicated by an upward shift of fluorescence and Coomassie-stained bands with increasing Frag D. Unbound Frag D appeared at ~8-fold molar excess (lane 7), suggesting a binding stoichiometry of ~7.

**Figure 5:**
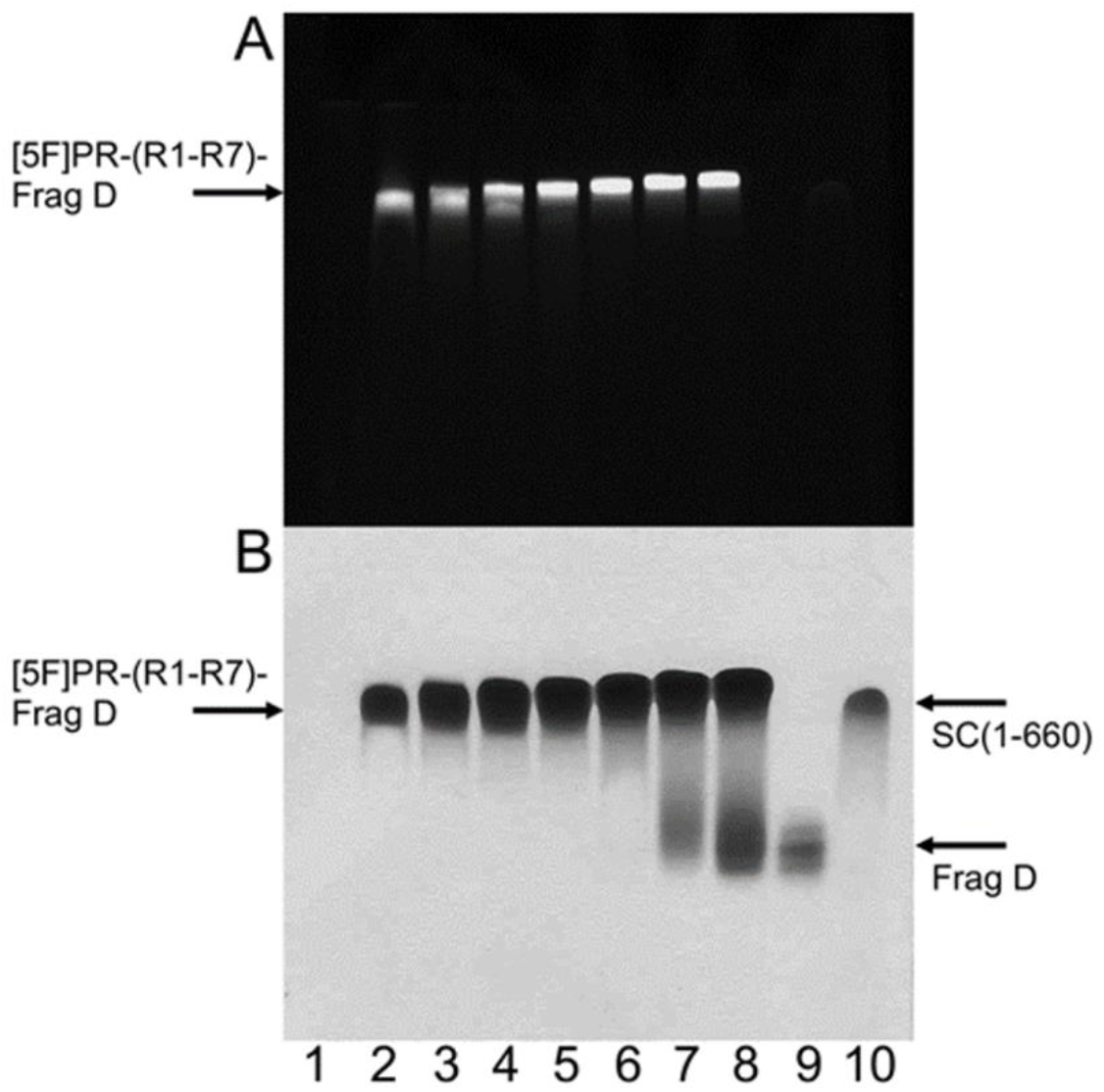
Binding of Frag D to [5F]PR-(R1→R7): ***A***, fluorescence and ****B****, Coomassie stain of incubations of [5F]PR-(R1→R7) (4.3 μM) with Frag D (lanes 2-8, respectively 10.0, 15.0, 20.0, 25.0, 30.0, 35.0 and 40.0 μM) at 25 °C for 30 minutes. Lanes 1, 9 and 10 are respectively [5F]PR-(R1→R7) (eluted with dye front and not visible), Frag D control and SC(1-660) as external control. Proteins were run on a 6 % Tris-Glycine native gel at 4 °C.

Frag D bound to [5F]PR-R1R6R7 and [5F]PR-R1R2R3, illustrating the presence of binding sites by continuous and non-continuous repeat sequences, independent of the order of the repeats. (**Fig. S3**). Unbound Frag D was observed above 3.5-fold molar excess, suggesting binding of ~3 Frag D molecules per construct, consistent with binding sites formed at the PR-R and R-R junctions.

### Equilibrium binding of [5F]-labeled C-terminal repeat peptides to Frag D

No increase in fluorescence intensity was observed when [5F]PR and [5F]R1 were titrated with Frag D, suggesting that these peptides do not bind Frag D (**Fig. 6A**), consistent with our native PAGE findings. In contrast, [5F]PR-R1, [5F]PR-R2, [5F]PR-R3, [5F]PR-R6 and [5F]PR-R7 bound to Frag D, with 1:1 stoichiometry and dissociation constants (*K*_D_) ranging from ~49 to ~130 nM (**Figs. 6B-F**). The results for [5F]PR-R1 were in excellent agreement with our previously reported *K*_D_ of 36 ± 8 nM for Frag D binding (15). The *K*_D_ values, *ΔF*_max_/*F*_o_ and stoichiometries obtained by quadratic fit, are listed in **Table 1**. Errors are given by 2x SD (95% confidence interval). The results indicate the presence of one single binding site for Frag D that bridges PR and the individual R repeats. Because R4 and R5 are identical to R2, their PR-R constructs were not tested for Frag D binding. Frag D is a dimer of three polypeptide chains, α, β, γ. The β and γ chains contain holes at their C-terminal end. The peptide Gly-Pro-Arg-Pro (GPRP) is known to bind and block the β- and γ-holes of Fbg and Frag D (31–33). [5F]PR-R1 was titrated with Frag D at saturating GPRP. In presence of 5 mM GPRP, [5F]PR-R1 still bound Frag D with *K*_D_ of 18 ± 10 nM, and *ΔF*_max_/*F*_o_ 0.08 ± 0.01 (**Fig. 7**), indicating that [5F]PR-R1, and most likely also the other PR-R constructs do not bind the β- and γ-chain holes of Frag D, but utilize a different binding site on Frag D, and that the SC C-terminal domain does not interfere with fibrin polymerization.

**Table 1:** Parameters of [5F]PR, [5F]R1, and [5F]PR-R binding to Frag D. Dissociation constants (*KD*), stoichiometric factors (*n*), and maximum fluorescence intensity (*ΔF_max_/F_o_*) were obtained by fitting of the titration data by the quadratic binding equation. Experimental errors represent ± 2 S.D. Equilibrium binding was performed as described under “Experimental Procedures.”

| | Stoichiometric factor<br>($n$) | $K_D$<br>($nM$) | $\Delta F_{max}/F_o$ |
| --- | --- | --- | --- |
| Probe | ($mol\ Frag\ D/mol\ repeat$) | | |
| [5F]PR | no binding |  |  |
| [5F]R1 | no binding |  |  |
| [5F]PR-R1 | $1.0 \pm 0.1$ | $49 \pm 14$ | $0.11 \pm 0.01$ |
| [5F]PR-R2 | $0.9 \pm 0.2$ | $130 \pm 42$ | $0.06 \pm 0.01$ |
| [5F]PR-R3 | $1.1 \pm 0.2$ | $99 \pm 26$ | $0.10 \pm 0.01$ |
| [5F]PR-R6 | $0.8 \pm 0.2$ | $115 \pm 47$ | $0.06 \pm 0.01$ |
| [5F]PR-R7 | $0.8 \pm 0.2$ | $112 \pm 25$ | $0.08 \pm 0.01$ |

**Figure 6:**
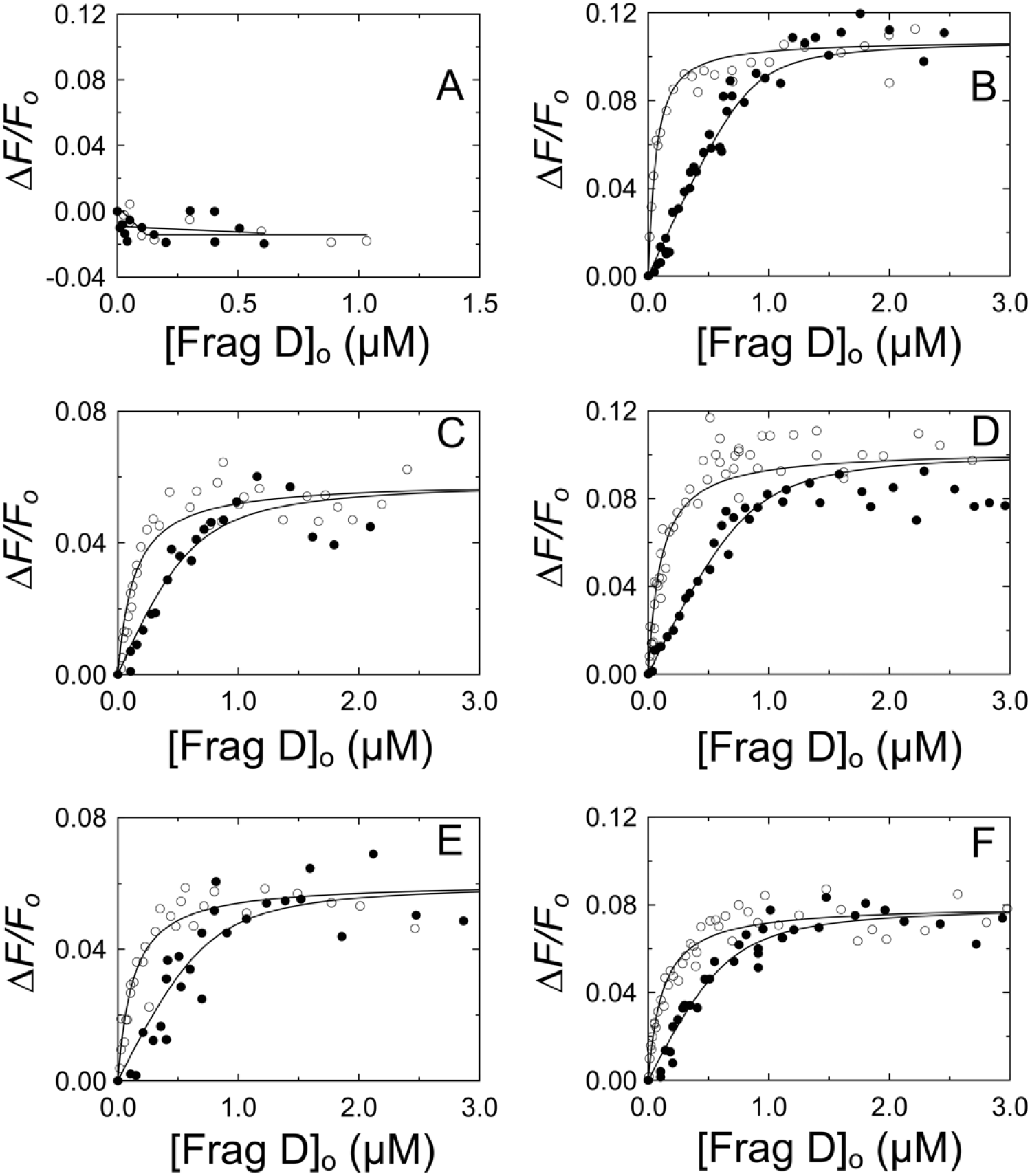
Fluorescence equilibrium binding of repeat constructs to Frag D. ***A,*** Titrations of 101 nM [5F]PR (○) and 67 nM [5F]R1 (●) with Frag D. ****B,**** Titrations of 25 (○) and 757 (●) nM [5F]PR-R1 with Frag D. ****C,**** Titrations of 9 (○) and 690 (●) nM [5F]PR-R2 with Frag D. ****D,**** Titrations of 9 (○) and 713 (●) nM [5F]PR-R3 with Frag D. ****E,**** Titrations of 21 (○) and 908 (●) nM [5F]PR-R6 with Frag D. ****F,**** Titrations of 15 (○), 39 (●) and 848 (Δ) nM [5F]PR-R7 with Frag D. *Solid black lines* represent the quadratic fit. Titrations and data fitting were performed as described in “Experimental Procedures.” The parameters are given in Table 1.

**Figure 7:**
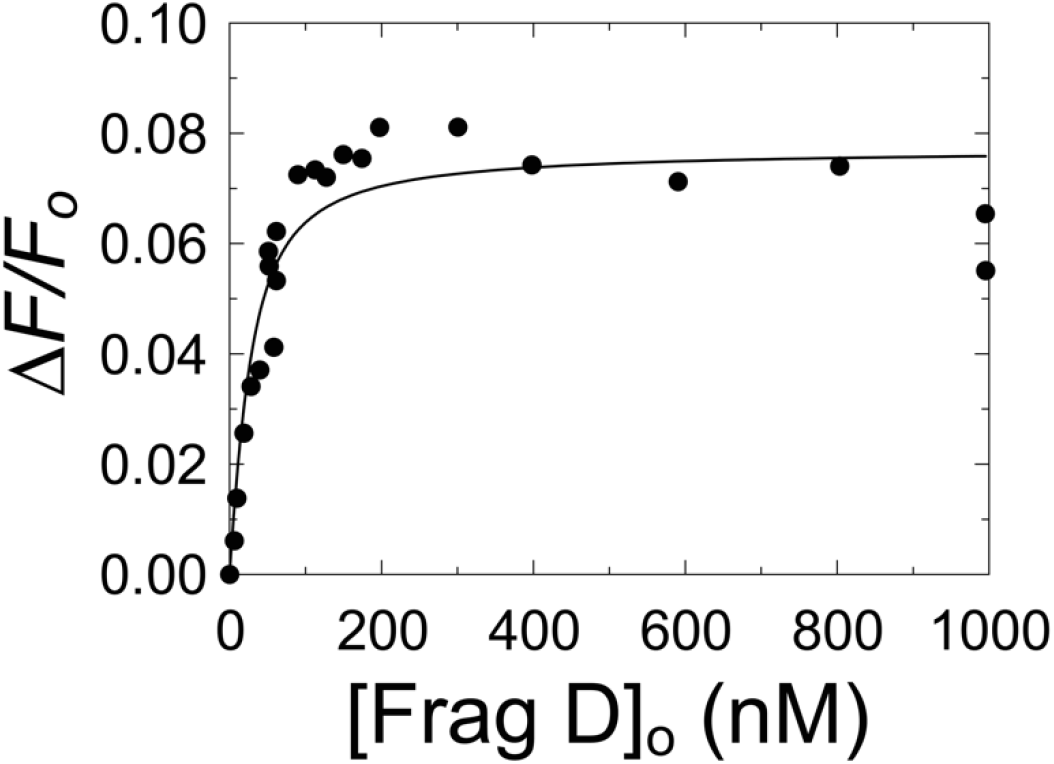
Fluorescence equilibrium binding of [5F]PR-R1 to Frag D in the presence of Gly-Pro-Arg-Pro (GPRP). The change in fluorescence intensity (*ΔF/F_o_*) of 19 nM [5F]PR-R1 as a function of total Frag D concentration ([Frag D]_o_). The *solid black line* represents the quadratic fit. Titrations and data fitting were performed as described in “Experimental Procedures.”

### NMR assignment of PR-R7, and titration of PR-R7 with Frag D

The ^1^H, ^15^N heteronuclear single quantum coherence (HSQC) spectrum and assignments of PR-R7, and its carbon direct detect spectra with full assignment of all residues were recently published (27), and chemical shift data are retrievable from the BMRB databank (http://www.bmrb.wisc.edu) under accession number 27036. The assignment for each correlation peak in the ^1^H-^15^N HSQC spectrum provides residue-specific information on an atomic level for each amide in the peptide. This allows observation of residue-specific changes of PR-R7 exposed to Frag D and results in a clearly defined interaction map. Full assignment was achieved with amide based experiments, supplemented with novel ^13^C-direct detect methods, allowing for full N, HN, CA and C resonance assignments, with the exception of the N- and C-terminal one (27). The ^13^C direct detect experiments unambiguously assigned residues that were problematic in amide based experiments: prolines; Glu^2^, Thr^3^, Tyr^24^, Tyr^34^, Arg^44^, and Val^72^ due to peak overlap; low intensity peaks for His^8^, Asn^16^, Asn^54^, His^61^, Ala^62^, and Ser^70^; and Tyr^9^ as it was between the unassigned His^8^ and Pro^10^ (**Fig. S4**). Peak overlap and low intensity peaks remained an issue for the analysis of the HSQC based titration experiments, but unambiguously knowing the exact position of these peaks from the ^13^C direct detect experiments was crucial during the refinement of the titration analysis. Given the very limited chemical shift dispersion of 8.2 ± 0.3 ppm in the proton dimension, which led to the peak overlap, it was anticipated that the PR-R7 peptide had limited to no secondary structure. With the backbone assignment in hand, secondary structure prediction using the program Talos + (34) confirmed that there was no consistent secondary structure to be anticipated in PR-R7 alone (**Fig. S5**).

With the HSQC spectrum assigned, binding of Frag D to PR-R7 was investigated in sodium phosphate buffer pH 7.0. The initial spectrum contained 65 μM of ^15^N-labeled PR-R7 peptide alone and exhibited all expected nitrogen-proton resonances. Upon successive addition of 240 μM aliquots of non-labeled Frag D, most resonances became weaker as more Frag D was added (**Fig. 8**). Individual titration points were chosen at 0.50, 1.00, 2.00, 3.21, and 4.95 mol-equivalents (me) of Frag D and PR-R7 peptide. Decreasing intensity of most resonances in the HSQC spectrum was attributed to intermediate exchange, and formation of a complex, affecting the correlation time of the molecule substantially broadening those resonances. The titration was analyzed by measuring the peak intensity for each residue of the first three titration spectra, recording them as a percentage of the reference spectrum (**Fig. 9**). The peak intensity change follows the same trend for each individual residue after each addition of Frag Only the first three titration points at 0.50, 1.00, and 2.00 me of Frag D contained meaningful information for the majority of residues and were analyzed. At higher Frag D concentrations most resonances were exchange broadened or affected by the slower tumbling of the larger Frag D·PR-R7 complex to the point where individual peaks were no longer contributing in a relevant matter to the analysis. Only about a dozen high intensity peaks remained visible in the spectra with 3.21 and 4.95 me of Frag D. Manual analysis of overlapping peaks assured correct intensity representation. HSQC spectra of residue pairs Glu^2^-Tyr^9^, Ser^7^-Thr^53^, Lys^20^-Asp^26^, Tyr^34^-Phe^39^, and Glu^33^-His^61^ were overlapping, with mutual influencing of peak intensities, which complicated independent analysis. The Glu^2^-Tyr^9^ and Glu^33^-His^61^ pairs merited further analysis. Glu peaks, typically weak to begin with, are often exchange-broadened in these spectra. The normally stronger Tyr^9^ was weakened by Frag D binding, making it look similar to the Glu^2^ peak. Similarly, Glu^33^ was additionally broadened by its position in the binding interface, while the overlapping His^61^ was exchange-broadened from the beginning, and this effect is accentuated in this environment, causing the intensities of these two peaks to decrease at similar rates. **Fig. 9** shows that the intensity of Glu^2^ and His^61^ is clearly smaller than anticipated compared to their neighbors, while Tyr^9^ and Glu^33^ fit in well with their neighbors, indicating that the intensity fit indeed was correct for those residues, but for different reasons. The intensities of other overlapping peaks matched well with those of their neighbors and had no impact on the overall interpretation. The correlation peaks in the HSQC spectrum at 2.0 me of Frag D for Lys^17^, Thr^29^, and Asn^35^ were no longer detectable, but the first two titration points showed clear signal intensities that were analyzed. More interestingly, **Fig. 9** reveals a trend where the stretch of residues 20 to 49 shows consistently more diminished signal intensity in all three titration points, compared to the C-terminal residues 51 to 75 and N-terminal residues 2 to 18 that retain a stronger intensity on average. The N-terminal residues 3 to 7 are affected the least and their signals remain visible even at the highest Frag D addition. The most plausible explanation is that Frag D has its main binding interaction with PR-R7 within the PR-R7 residues 20 to 49, which suggests a binding site overlapping PR and R7. Significantly less signal reduction was observed for the peaks in the C-terminal residues 51 to 75 leading to the conclusion that this portion of the peptide might contribute less to the interaction with Frag D, although weak binding of that sequence with Frag D can still be anticipated. The stretch of residues from 22 to 50 was determined to constitute the minimal peptide (MP) that is required for binding to Frag D. Because the NMR studies required 20 mM sodium phosphate, 150 mM NaCl, pH 7.0 buffer, equilibrium binding of Frag D to [5F]PR-R7 was also performed in phosphate buffers with corresponding ionic strength at pH 7.0 and 7.4 to determine whether the phosphate buffer and pH affect binding. Titration of [5F]PR-R7 with stoichiometries fixed at 1, yielded *K*_D_ values of 3 ± 2 μM, and 27 ± 75 nM at pH 7.0 and 7.4, respectively. The data at pH 7.4 corresponded well with the results for Frag D binding in HEPES buffer. Titrations in phosphate buffer were somewhat noisier than in HEPES buffer, presumably due to probe-buffer interactions. Binding at pH 7.0 was weaker, but consistent with the NMR binding data collected at micromolar reactant concentrations.

**Figure 8:**
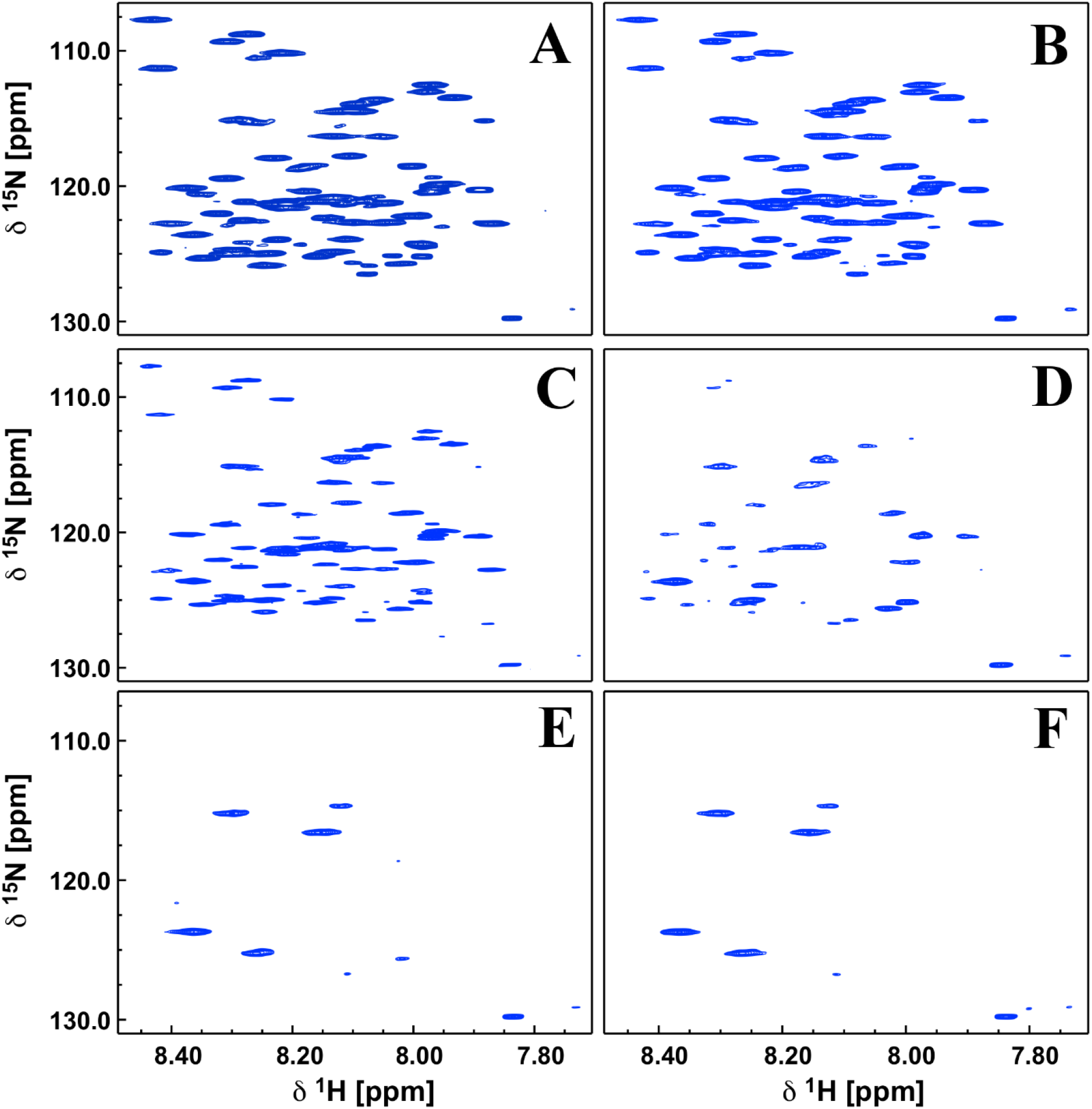
NMR titration of PR-R7 with Frag D. The HSQC spectra of ^15^N labeled PR-R7 peptide were collected at 600MHz and 298K. Increasing amounts of unlabeled Frag D were added and a HSQC was measured with the same parameters, yielding spectra ****A**** through ****F**** representing the reference spectrum, with no Frag D added in ****A****, and 0.5, 1.0, 2.0, 3.21, and 4.95 me of Frag D present in spectra ****B**** through ****F****. The spectra clearly show the diminishing peak intensity upon increasing Frag D addition, indicating intermediate exchange and complex formation with Frag D.

**Figure 9:**
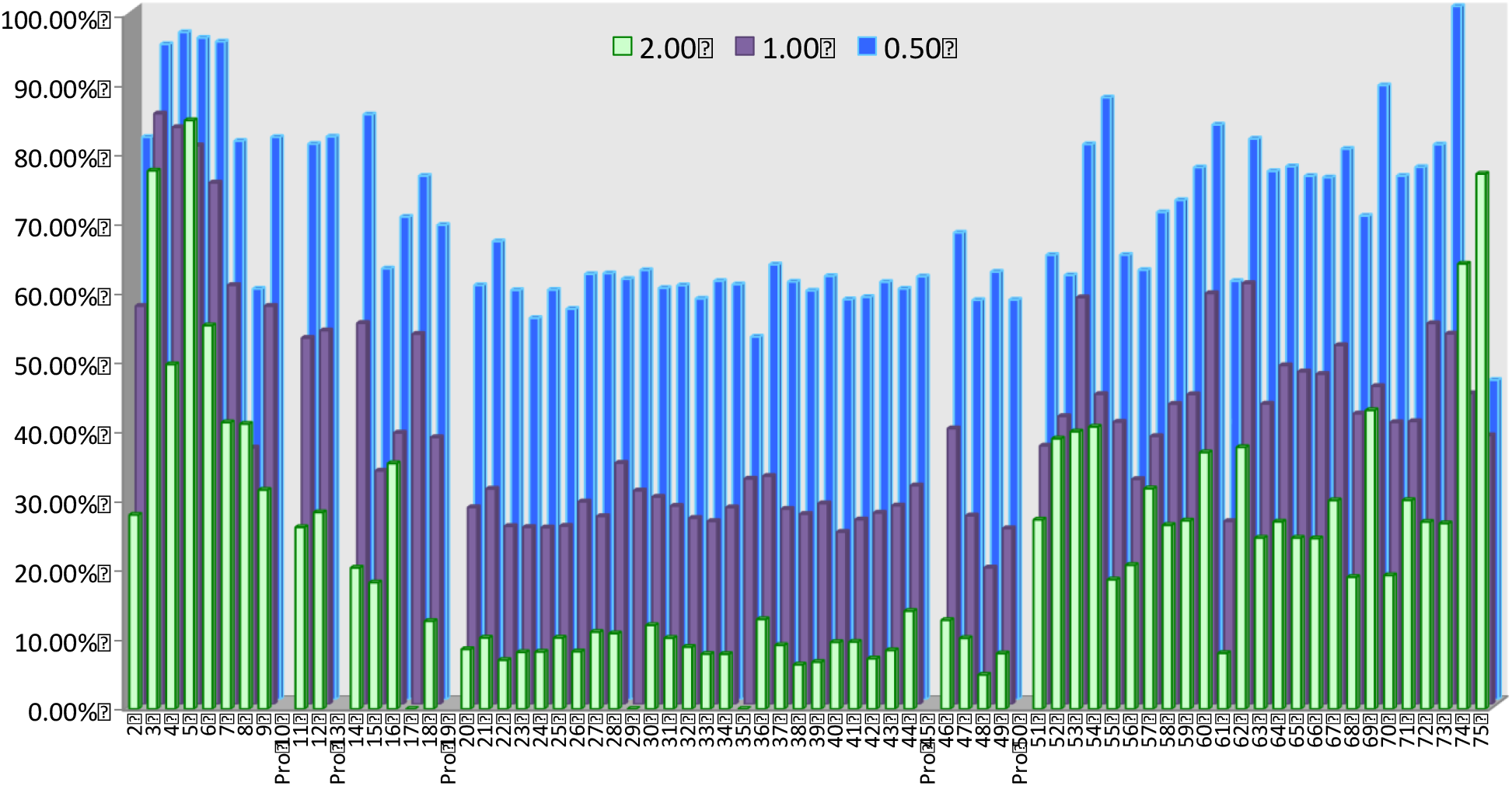
Residue-specific response of PR-R7 in the NMR titration with Frag D. *Blue bars* represent the residual percent intensity of the HSQC correlation peak for the respective residue at 0.5 molar equivalent Frag D; *purple* at 1 molar equivalent and *green* at 2 molar equivalent. The intensity is based on the reference spectrum of 65 μM of ^15^N-labeled PR-R7 peptide alone. Aliquots of 240 μM Frag D stock solution were added to achieve the 0.5, 1.0, and 2.0 me conditions.

### Equilibrium binding of Frag D to the 5-fluorescein-labeled Minimal Peptide, [5F]MP

The 29-residue MP, identified by NMR studies, was synthesized, with an additional C-terminal Cys for attachment of a 5-fluorescein probe. Titration of [5F]MP (23 nM) with Frag D demonstrated binding, with *K*_D_ of 5 ± 4 μM, *ΔF_max_/F_o_* 0.17 ± 0.02, and a fixed stoichiometry of 1 (**Fig. 10**). This indicates that the MP sequence is sufficient to bind Frag D.

**Figure 10:**
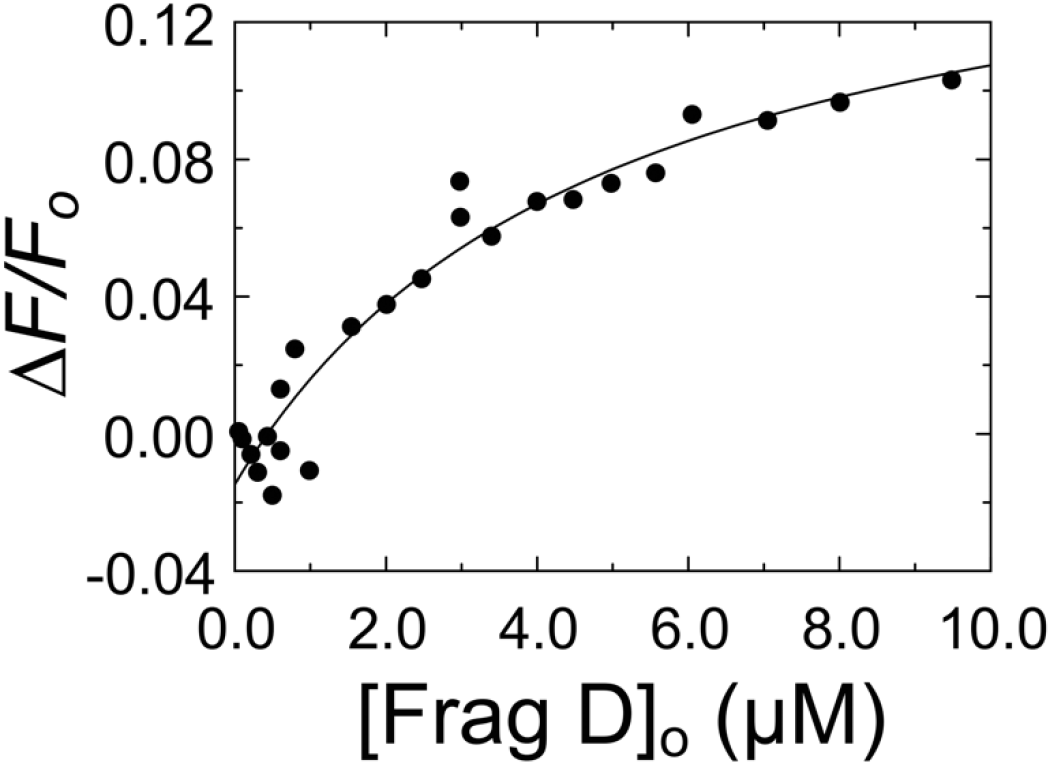
Fluorescence intensity titration of [5F]MP with Frag D. Change in fluorescence intensity (*ΔF/F_o_*) of 23 nM [5F]MP as a function of total Frag D concentration ([Frag D]_o_). The *solid black line* represents the quadratic fit. Titrations and data fitting were performed as described in “Experimental Procedures.”

### Equilibrium binding of alanine scanning mutants of PR-R7 to Frag D

Alanine scanning mutagenesis and equilibrium binding further determined the specific PR-R7 residues involved in Frag D binding. Sequential mutation of 4-6 amino acid residue stretches was done in the MP region and beyond, until the end of repeat R7. Ten alanine mutant constructs of [5F]PR-R7 were titrated with Frag D. Alanine mutants of VKYRDA, GTGIR, EYNDG in PR, and of ARPT, YKKP, and HADG in R7 abolished binding (**Figs. 11, 12A**), indicating essential contributions of these sequences to Frag D binding. VKYRDA, GTGIR and EYNDG form a contiguous stretch located at the C-terminal region of the PR. ARPT and YKKP are located at the N-terminal region of R7 whereas HADG is located toward the C-terminus of R7. Mutations at TFGYE, SETNA, YNVTT and TATYG still allowed binding of Frag D, with respective *K*_D_ values of 500 ± 400 nM, 120 ± 40 nM, 350 ± 150 nM, and 260 ± 100 nM (**Figs. 11, 12B-E**). Alanine substitution at SETNA resulted in a binding affinity similar to that of [5F]PR-R7, suggesting limited involvement of these residues in binding. Maximum fluorescence intensity increases of these 4 constructs were comparable to that of native [5F]PR-R7. The sequence alignment of the corresponding MP regions in other inter-repeat regions showed strictly conserved G^37^, Y^41^, A^43^, R^44^, P^45^, K^49^ and P^50^ (**Fig. 13**). G^28^ and I^31^ are present only in the PR, and the corresponding residues A and V in other repeats are hydrophobic and similar in size. Y^41^ is highly conserved, however Ala substitution still allows for weak binding of Frag D.

**Figure 11:**
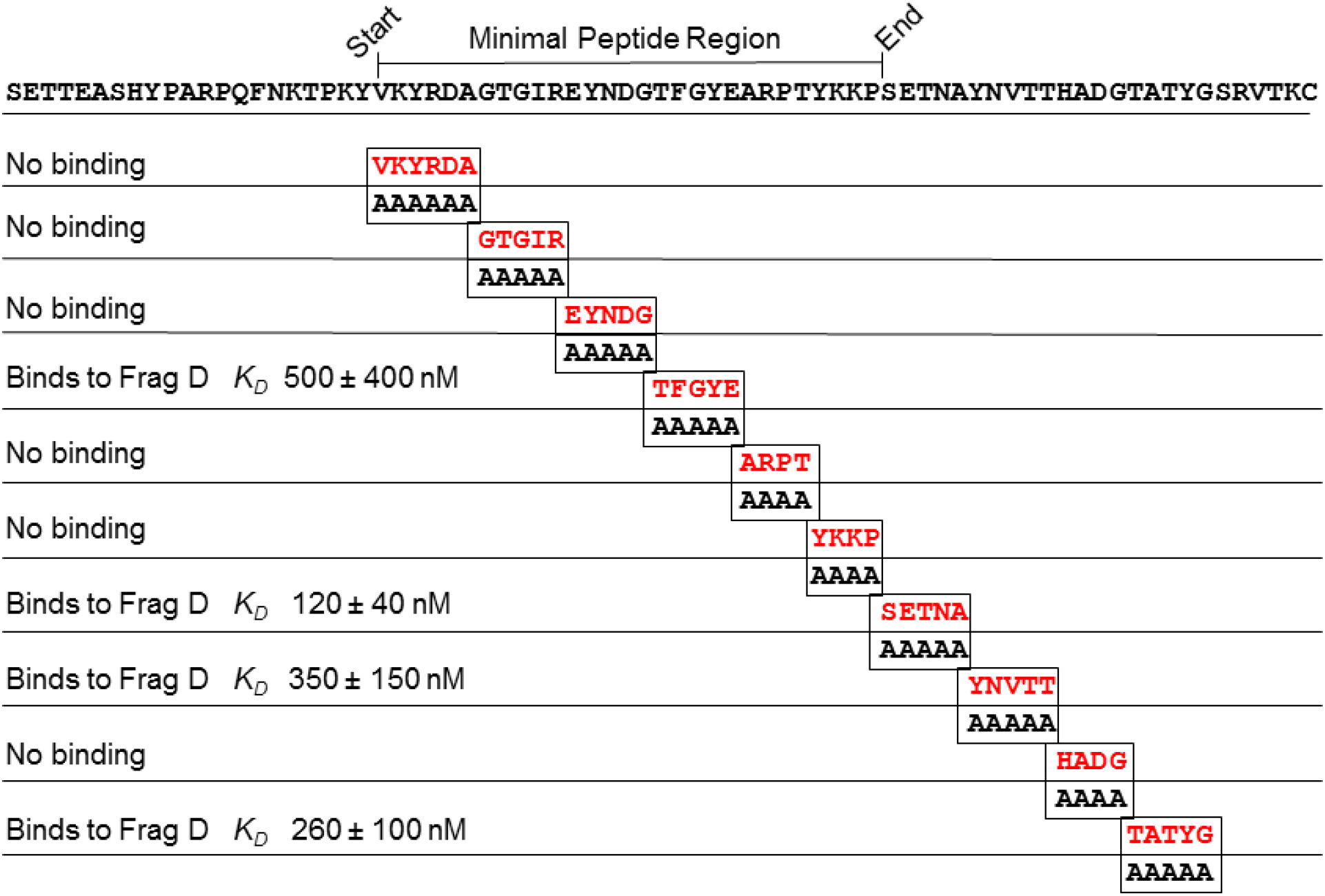
Binding of Frag D to PR-R7 alanine mutants. Alanine scanning mutagenesis of the MP region and the remainder of the R7 C-terminal region. The MP consists of a continuous stretch of 29 residues of which 21 are in the PR, and 8 in R7.

**Figure 12:**
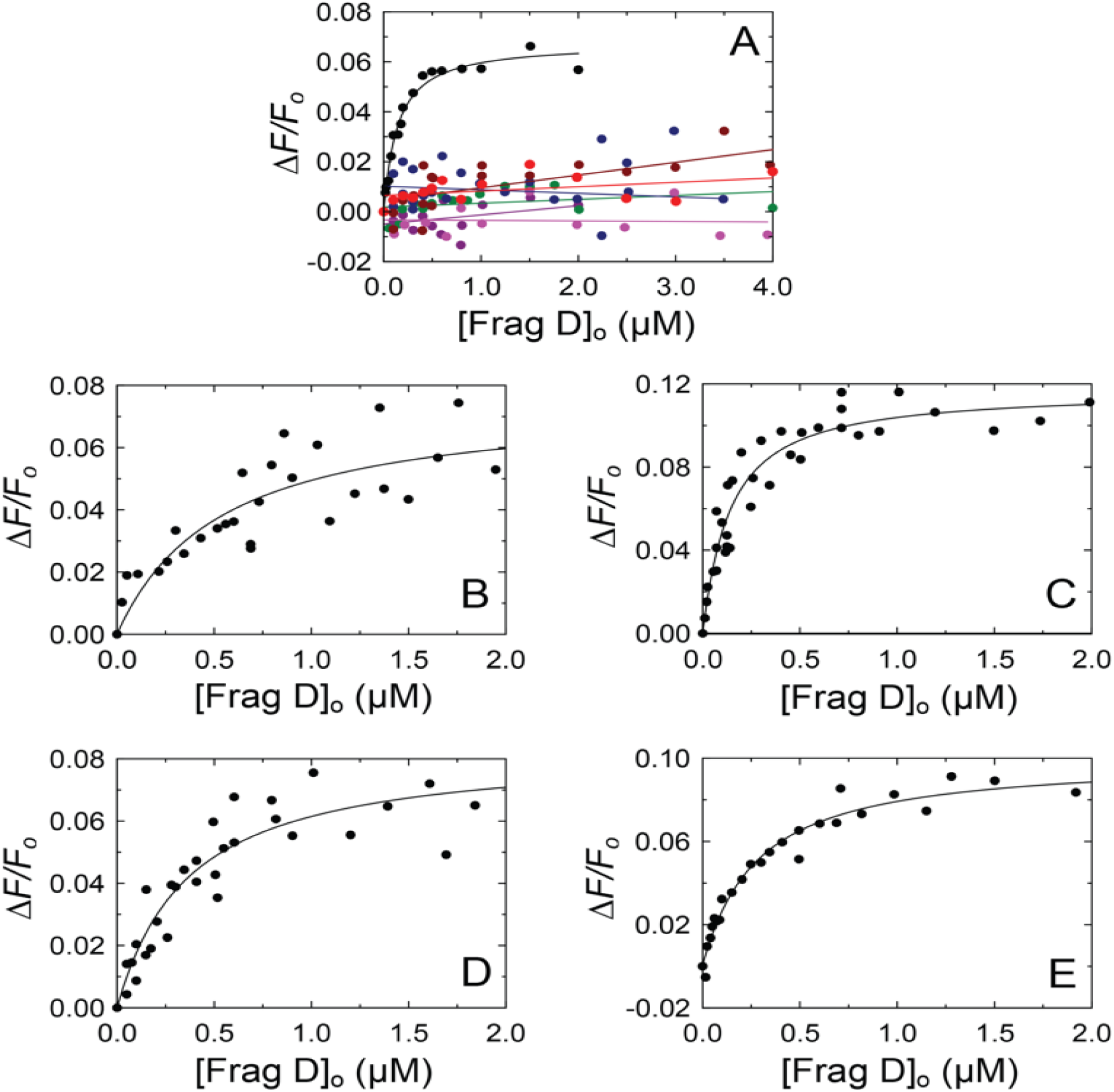
Equilibrium binding of Frag D to [5F]PR-R7 alanine mutants. ***A***. The change in fluorescence intensity (*ΔF/F_o_*) of 19 nM [5F]PR-R7 (●), 33 nM [5F]PR-R7-VKYRDA (●), 39 nM [5F]PR-R7-GTGIR (●), 35 nM [5F]PR-R7-EYNDG (●), 18 nM [5F]PR-R7-ARPT (●), 24 nM [5F]PR-R7-YKKP (●) and 29 nM [5F]PR-R7-HADG (●) as a function of total Frag D concentration ([Frag D]_o_). ****B****. *ΔF/F_o_* of 37 nM [5F]PR-R7-TFGYE as a function of [Frag D]_o_. ****C****. *ΔF/F_o_* of 27 nM [5F]PR-R7-SETNA as a function of [Frag D]_o_. ****D****. *ΔF/F_o_* of 27 nM [5F]PR-R7-YNVTT as a function of [Frag D]_o_. ****E****. *ΔF/F_o_* of 27 nM [5F]PR-R7-TATYG as a function of [Frag D]_o_. *Solid lines* represent the quadratic fit. Titrations and data fitting were performed as described in “Experimental Procedures.”

### Clotting of Fbg by the SC(1-325)·ProT^QQQ*^ and SC(1-660)·ProT^QQQ*^ complexes

Cleavage of Fbg to Fbn causes turbidity increase due to formation and polymerization of Fbn with a lower solubility. We compared the rates of Fbn clotting by thrombin, and the active SC(1-325)·ProT^QQQ*^ and SC(1-660)·ProT^QQQ*^ complexes. ProT(R155Q,R271Q,R284Q), or ProT^QQQ^, is a human ProT variant with its prothrombinase and thrombin cleavage sites mutated to prevent proteolytic processing (35). It can be activated proteolytically to a meizothrombin form, as well as conformationally by triggering formation of an active site by occupation of its Ile^16^ pocket by the N-terminus of SC. Thrombin, as a control reaction, initiated immediate cleavage of Fbg whereas addition of ProT^QQQ^ in mixtures with SC(1-325) and SC(1-660) showed a lag phase during which the SC(1-325)·ProT^QQQ*^ and SC(1-660)·ProT^QQQ*^ complexes are formed (**Fig. 14**). In the thrombin- or SC(1-325)·ProT^QQQ*^-initiated cleavage of Fbg at 0.5 mg/ml there was rapid and comparable increase in turbidity reaching a maximal level, indicating clot stabilization. Clot formation by SC(1-660)·ProT^QQQ*^ was slower, and the initial rates of turbidity increase, and absorbance readings at clot stabilization were dependent on the Fbg concentration. SC(1-325) conformationally activates ProT but lacks the C-terminal domain for Frag D binding. This additional interaction of SC(1-660) may sequester Fbg molecules at the C-terminal domain, thereby decreasing the effective *in vitro* Fbg concentration available for cleavage by SC(1-660)·ProT^QQQ*^, or posing conformational restrains on cleavage of Fbg bound in the substrate mode, either with or without simultaneous tethering to the SC repeat domain.

**Figure 13:**
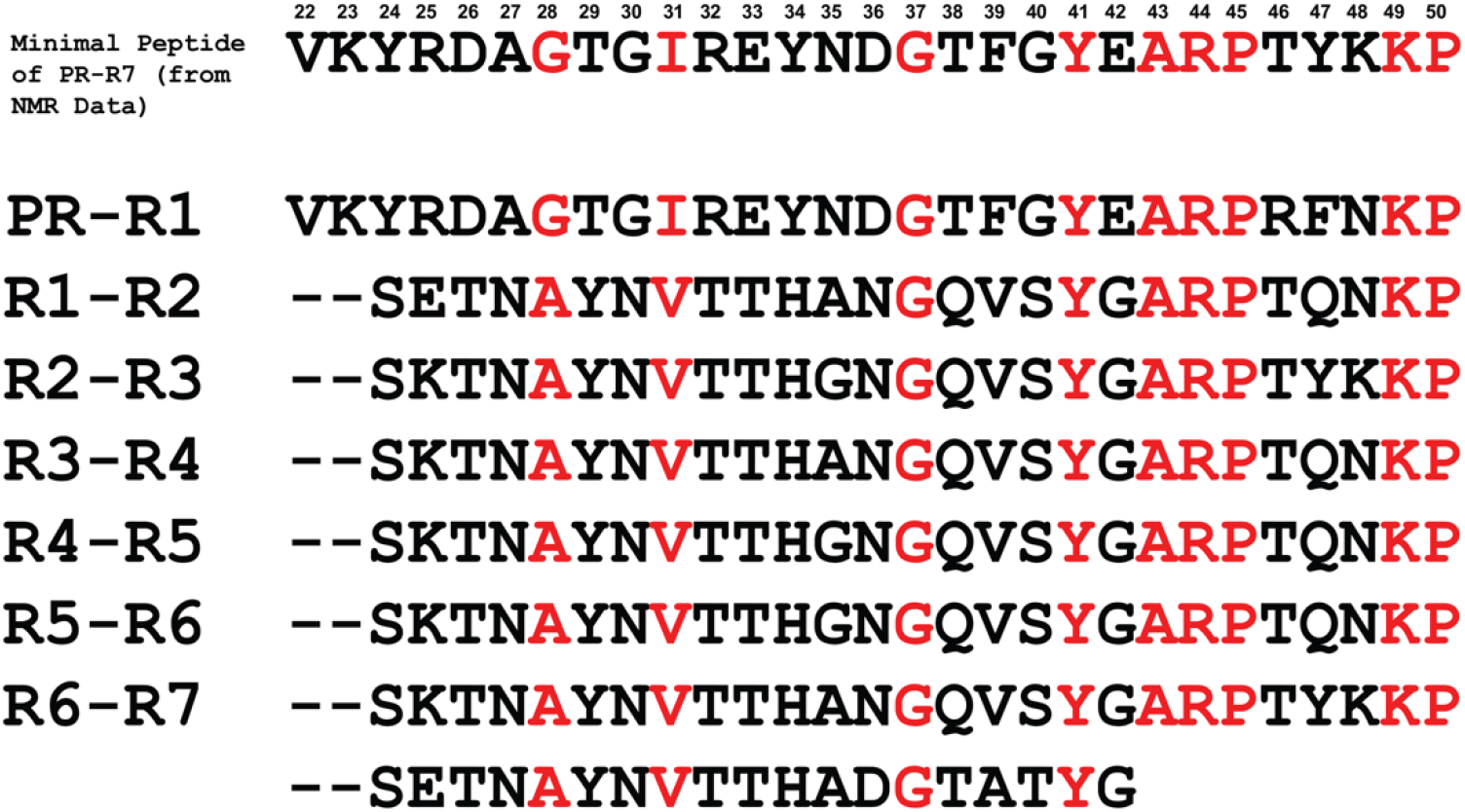
MP sequence alignment with the bridging sequences between SC repeats. The alignment shows the conserved residues (*red*) at positions G^37^, Y^41^, A^43^, R^44^, P^45^, K^49^ and P^50^. The residues G^28^ and I^31^ are present only in the PR, and are replaced with similar size hydrophobic residues A and V in other repeats. Residue numbering is consistent with the PR-R7 sequence.

**Figure 14:**
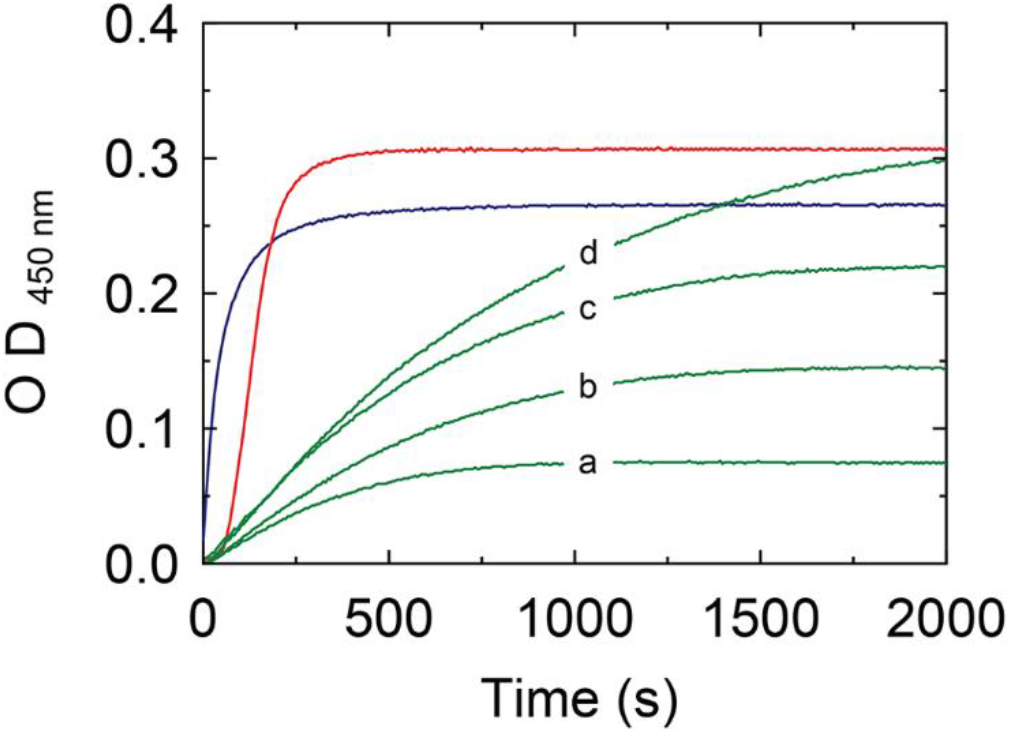
Turbidity assay of Fbg cleavage: Increase in turbidity at 450 nm, 25 °C for mixtures of 10 nM thrombin with 0.5 mg/ml Fbg (*blue*); 10 nM SC(1-325)·ProT^QQQ*^ complex with 0.5 mg/ml Fbg (*red*); and 10 nM SC(1-660)·ProT^QQQ*^ complex with increasing Fbg (*green*, *a-d*; 0.3, 0.5, 0.75 and 1.0 mg/ml respectively).

### Fluorescence anisotropy titrations of repeat constructs binding to Frag D

The full-length repeat construct, PR-(R1→R7), contains the highly conserved PR and 7 repeats, of which R2, R4 and R5 are identical. Constructs for direct or competitive binding, compared to full-length SC(1-660), are shown in **Fig. 1**. Using fluorescence anisotropy, we titrated three independently prepared batches of [5F]PR-(R1→R7) (2 concentrations, given in figure legend) with Frag D (**Fig. 15, A1,B1,C1**). Their [5F]PR-(R1→R7)·Frag D complexes (2 concentrations) were used in competitive titrations with three independently prepared PR-R1R2R3 batches (**Fig. 15, A2, B2, C2**) to determine *K*_D_ values for PR-R1R2R3 and to validate the assay and reproducibility. **Fig. 15, B3, C3, C4** show competitive titrations of the [5F]PR-(R1→R7)·Frag D complex (2 concentrations) with PR-R1R6R7, PR-R3R4R7 and PR-R3R6R7 measured by fluorescence anisotropy. Inter-batch results were in good agreement, with stoichiometries of ~5 for Frag D binding to [5F]PR-(R1→R7), *K*_D_s of ~7 - 32 nM, maximum anisotropy changes *r*_max_ 0.13 - 0.15, and *r*_0_ initial anisotropy ~0.1 (**Table 2**). Competitive binding of Frag D to PR-R1R2R3, PR-R1R6R7, PR-R3R4R7 and PR-R3R6R7 gave stoichiometries of ~3 as expected for a construct containing 3 inter-repeat sequences with conserved residues essential for Frag D binding, and binding parameters comparable to those of [5F]PR-(R1→R7) (**Table 2**). Repeat swapping maintained these conserved inter-repeat sequences and did not weaken the binding of Frag D significantly.

**Table 2:** Binding parameters from titrations of repeat constructs with Frag D. Three independent [5F]PR-(R1→R7) preparations were titrated with Frag D, and the dissociation constant (*K*_D_), stoichiometric factor (*n*), initial anisotropy (*r_o_*), and maximum anisotropy (*r_max_*) were obtained by fitting to the quadratic binding equation. Competitive binding of PR-R1R2R3 (three independent preparations), PR-R1R6R7, PR-R3R4R7, and PR-R3R6R7 to Frag D was measured by titration of two fixed [5F]PR-(R1→R7)/Frag D mixtures with the unlabeled peptides. The two curves of each data set were fit simultaneously with the two direct [5F]PR-(R1→R7) binding curves, using the cubic equation, to obtain *K*_D_ and *n* for the competitor peptides. Experimental error represents ± 2 S.D. Binding studies and data analysis were performed as described under “Experimental Procedures.”

| Repeat Constructs | Stoichiometric factor<br>(mol Frag D/mol repeat) ($n$) | $K_D$<br>(nM) | $r_{max}$ | $r_o$ |
| --- | --- | --- | --- | --- |
| PR-R1R2R3 | $3.3 \pm 0.4$ | $25.0 \pm 8.0$ | $0.15 \pm 0.01$ | $0.10 \pm 0.01$ |
| | $3.5 \pm 0.8$ | $7.0 \pm 6.0$ | $0.13 \pm 0.01$ | $0.09 \pm 0.01$ |
| | $2.5 \pm 0.3$ | $7.0 \pm 3.0$ | $0.14 \pm 0.01$ | $0.10 \pm 0.01$ |
| PR-R1R6R7 | $3.0 \pm 0.7$ | $8.0 \pm 9.0$ | $0.13 \pm 0.01$ | $0.09 \pm 0.01$ |
| PR-R3R4R7 | $2.8 \pm 0.6$ | $42.0 \pm 16.0$ | $0.15 \pm 0.01$ | $0.10 \pm 0.01$ |
| PR-R3R6R7 | $3.0 \pm 0.8$ | $42.0 \pm 21.0$ | $0.15 \pm 0.01$ | $0.10 \pm 0.01$ |
| [5F]PR-(R1→R7) | $5.2 \pm 0.3$ | $12.0 \pm 5.0$ | $0.14 \pm 0.01$ | $0.10 \pm 0.01$ |
| | $4.8 \pm 0.6$ | $7.4 \pm 6.0$ | $0.13 \pm 0.01$ | $0.09 \pm 0.01$ |
| | $5.1 \pm 0.5$ | $32.0 \pm 9.0$ | $0.15 \pm 0.01$ | $0.10 \pm 0.01$ |

**Figure 15:**
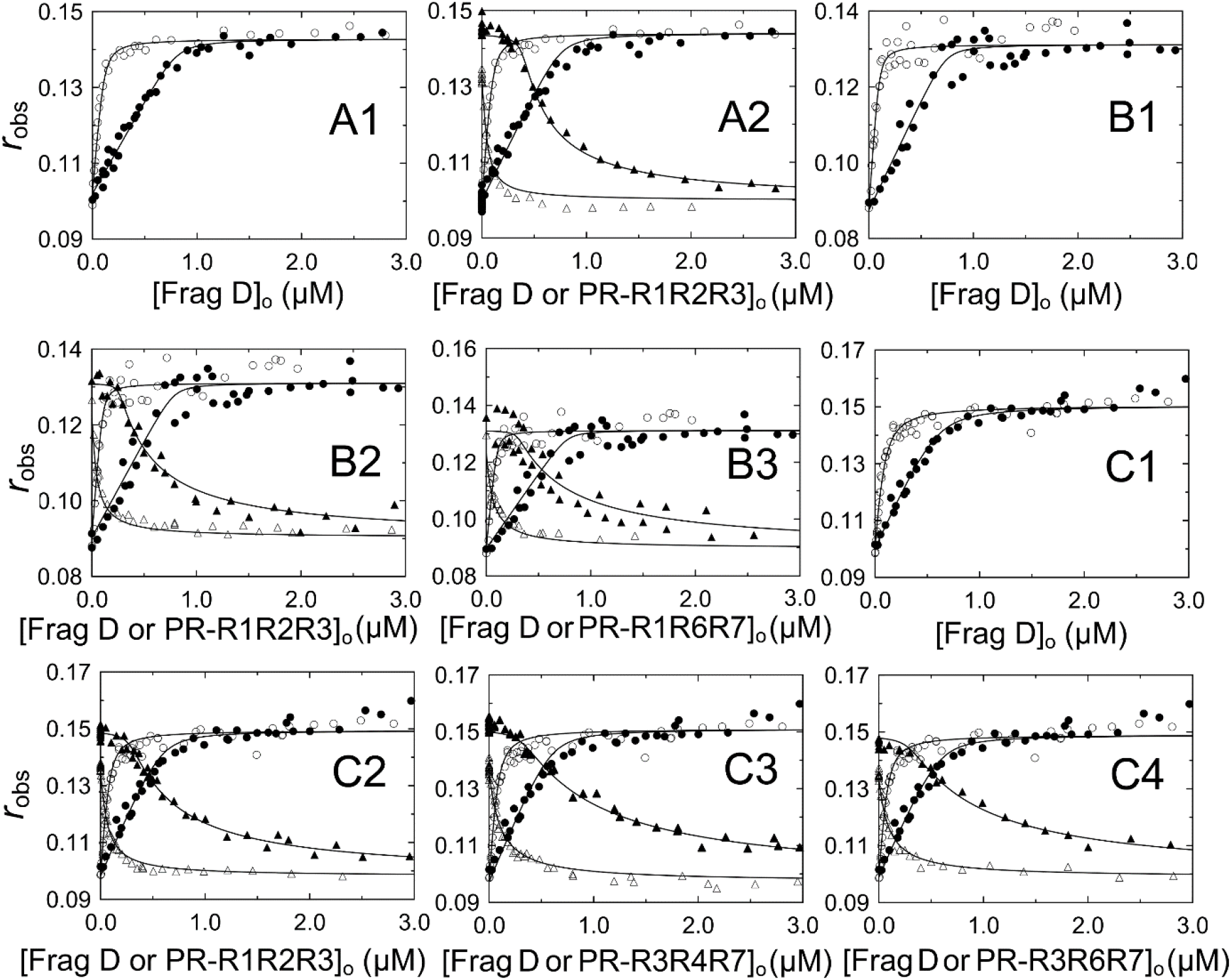
Fluorescence anisotropy titrations of repeat constructs binding to Frag D: ***A1***, observed anisotropy (*r_obs_*) of 21 (○) and 154 (●) nM [5F]PR-(R1→R7) (preparation 1) as a function of total Frag D concentration ([Frag D]_o_); ****A2****, simultaneous fit of ****A1**** with competitor PR-R1R2R3 (preparation 1) titrated into mixtures of 21 nM [5F]PR-(R1→R7) with 105 (**Δ**) and 1053 (**▲**) nM Frag D; ****B1****, *r_obs_* of 21 (○) and 155 (●) nM [5F]PR-(R1→R7) (preparation 2) as a function of total Frag D; ****B2****, simultaneous fit of ****B1**** with competitor PR-R1R2R3 (preparation 2) titrated into mixtures of 21 nM [5F]PR-(R1→R7) with 107 (**Δ**) and 1012 (**▲**) nM Frag D; ****B3****, simultaneous fit of ****B1**** with competitor PR-R1R6R7 titrated into mixtures of 21 nM [5F]PR-(R1→R7) with 107 (**Δ**) and 959 (**▲**) nM Frag D; ****C1****, *r_obs_* of 16 (○) and 120 (●) nM [5F]PR-(R1→R7) (preparation 3) as a function of total Frag D; ****C2****, simultaneous fit of ****C1**** with competitor PR-R1R2R3 (preparation 3) titrated into mixtures of 16 nM [5F]PR-(R1-R7) with 104 (**Δ**) and 1040 (**▲**) nM Frag D; ****C3****, simultaneous fit of C1 with competitor PR-R3R4R7 titrated into mixtures of 16 nM [5F]PR-(R1→R7) with 107 (**Δ**) and 1008 (**▲**) nM Frag D; ****C4****, simultaneous fit of ****C1**** with competitor PR-R3R6R7 titrated into mixtures of 16 nM [5F]PR-(R1→R7) with 99 (**Δ**) and 992 (**▲**) nM Frag D. Titrations and data analyses were performed as described under “Experimental Procedures.” *Solid black lines* represent the quadratic (****A1****, ****B1****, ****C1****) and cubic binding fits (all other panels). Binding parameters are given in Table 1.

### Circular dichroism spectroscopy of SC repeat peptides and Frag D

CD spectroscopy of full-length SC(1-660) (*blue*), SC(1-325) (*brown*), PR-R7 (*black*), PR-(R1→R7) (*red*) and Frag D (*green*) was performed to determine the presence of secondary structure content (**Fig. 16**). Frag D showed major helical content with typical minima at 208 and 222 nm. The negative band at 224 nm in the SC(1-325) spectrum was suggestive of helix (36), consistent with its known crystal structure (19). The profiles of PR-R7, PR-(R1→R7) and SC(1-660) indicated an extended or irregular structure. The 222/208 nm ratio of > 1.1 for Frag D suggested the presence of coiled-coil helices, consistent with its published structure (32). However, the PR-(R1→R7) spectrum was more ambiguous, with a positive band at 212 nm suggesting random coil (37,38). Although the 222/208 nm ratio was > 1, secondary structure predictions by GOR IV (39) and PSIPRED (40) gave ~70% random coil and ~30% extended strand content for PR-(R1-R7) and PR-R7, with no helical content. The respective helix, extended strand and random coil content estimates for SC(1-660) were 26, 20 and 54% (GOR IV) and 28, 12 and 60% (PSIPRED); and for SC(1-325) 50, 13 and 37% (GOR IV) and 54, 1 and 45% (PSIPRED). Prediction of natural disordered regions in full-length SC(1-660) by PONDR-FIT (41) indicated an overall disordered content of 46.21%, with 75.66% of the R1-R7 sequence identified as disordered.

**Figure 16:**
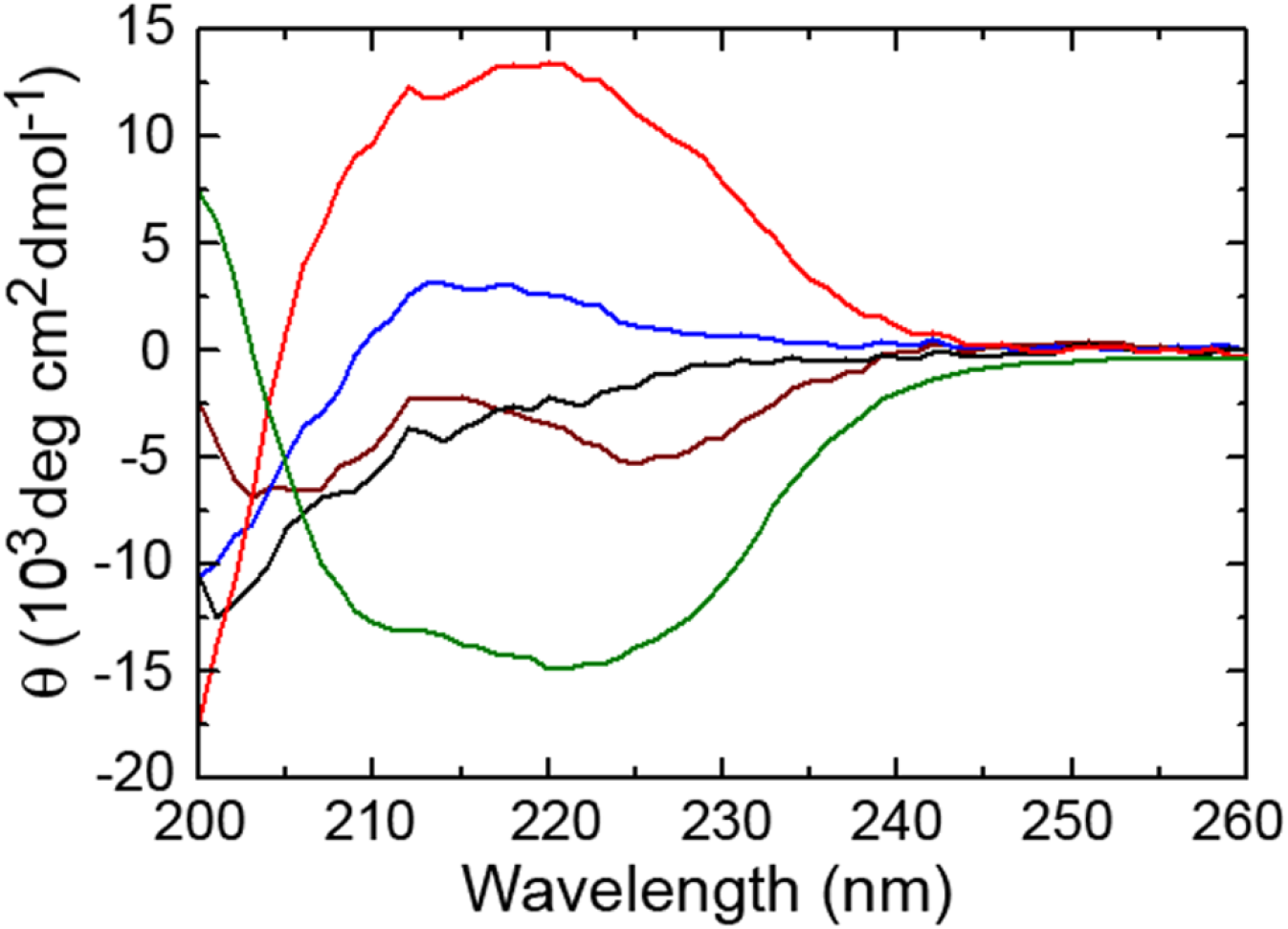
Circular Dichroism Spectra of Different SC Proteins and Frag D. CD spectra of PR-R7 (*black*), PR-(R1→R7) (*red*), SC(1-325) (*brown*), SC(1-660) (*blue*), and fragment D (*green*) were measured using a Jasco J-810 spectrometer at 25 °C.

## Discussion

Coagulase-positive *S. aureus*, a major threat to public health, employs various Fbg-binding virulence factors, including the blood clotting activator SC to facilitate rapid colonization and spreading inside the human host (42–47). We showed previously that host ProT is activated conformationally by insertion of the SC N-terminus in the Ile^16^-binding pocket, without the requirement for proteolytic activation (19,20). The SC·ProT* complex proteolytically converts substrate Fbg to fibrin clots that serve as focal points for bacterial adhesion (21). In addition, localization of SC on host fibrin(ogen) is facilitated by the SC C-terminal repeat sequence (15,28,29,48). SC serotypes have 4 to 9 repeats, preceded by a highly conserved PR containing a C-terminal TFGYE sequence not present in the repeats (3,22,49) (**Figs. 2, S1**). The presence of these highly conserved repeats in various *S. aureus* strains may be in part the result of selective pressure, increasing the Fbg-binding efficiency of SC. Our studies used SC of the prototypical Newman D2 Tager 104 strain that has 7 repeats exhibiting a high degree of conservation, with 100% identical R2, R4 and R5, suggesting duplication in the genome (19).

We used the Frag D domain of Fbg/fibrin, a ligand of choice, to demonstrate binding to free and ProT-complexed full-length SC(1-660), and C-terminal constructs containing PR and one or more repeats; the MP identified by NMR; and absence of binding to PR, the single repeats, or specific Ala-scanned PR-R7 constructs. The PR-R constructs with one repeat exhibited similar affinities for Frag D, suggesting a fairly conserved binding site, even with minor sequence variability among the repeats. We chose PR-R7, with the highest expression levels, for independent and mutually validating binding approaches because NMR typically requires fairly high experimental concentrations.

We identified the PR sequence VKYRDAGTGIREYNDG, strictly conserved among all serotypes, and ARPTYKKP and its respective similar sequences in the N-terminal portion of the repeats, as essential determinants in Frag D binding. Alignment of the minimal peptide MP with PR and the repeats shows 9 conserved residues (**Fig. 13**), G/A^28^, I/V^31^, G^37^, Y^41^, A^43^, R^44^, P^45^, K^49^, P^50^, of which G^37^, Y^41^, A^43^, R^44^ and P^45^ are at the PR-R and R-R junctions. Ala substitution of these residues abolishes or severely weakens Frag D binding, supporting Frag D-binding sites at the junctions of PR-R and R-R. The C-terminal TFGYE stretch in MP contains Y^41^ that is conserved in the inter-repeat domain sequences (**Fig. 13**). Ala substitution weakened Frag D binding ~5-fold, suggesting a role for the conserved Y in inter-repeat interactions with Frag D. HADG at the C-terminal end of PR-R7, aligns with YNDG in the MP, containing the conserved G^37^. Disruption of Frag D binding by Ala substitution of either sequence (**Figs. 11, 12A**) indicates that this G residue is a part of multiple Frag D/Fbg binding sites in the C-terminal repeat region. Our equilibrium binding and native PAGE data showed that PR and R1 by themselves do not bind Frag D with measurable affinity. However, all the PR-R constructs, containing the conserved G/A^28^, I/V^31^ and G^37^ residues aligned in the R-R bridging sequences, bound Frag D with a 1:1 stoichiometry and *K*_D_ values in the ~50 - 130 nM range. These findings, together with our Ala scanning results, are consistent with a simultaneous requirement for both binding subsites. The [5F]-labeled synthetic MP bound Frag D with *K*_D_ ~5 μM, and the tighter binding of PR-R (**Table I**) indicates that repetitive sequences formed by PR-R and R-R provide additional binding interactions for Frag D. Our findings support multiple binding sites for Frag D/Fbg, perhaps as many as 5-6, in the C-terminal domain of SC (15). The capacity to bind multiple Fbg/fibrin molecules simultaneously may facilitate bacterial localization on blood clots, evasion of the host immune system (50), and may impair the efficiency of antibiotic therapy.

N-terminal cleavage of the Fbg α- and β- chains to form fibrin monomer releases the fibrinopeptides AαBβ, with generation of new Gly-Pro-Arg N-termini that respectively bind the γ- and β-holes to form the fibrin network. The tetrapeptide GPRP binds to Fbg and Frag D holes with an affinity of ~5 mM (31,32), and is routinely used to prevent fibrin polymerization in solution studies of fibrin monomer. Blocking the Frag D holes with GPRP had no weakening effect on its binding to PR-R1 (**Fig. 7**), excluding them as potential interaction sites. This implies that interaction sites of Frag D are located elsewhere in the Frag D molecule, possibly in the coiled-coiled region. The modestly tighter affinity for Frag D binding (18 nM) compared to that in the absence of GPRP (~36 - 49 nM) may be the result of subtle conformational changes.

The NMR data further confirmed our hypothesis of a functional binding unit comprised of residues from both PR and the repeats. Decreased NMR peak intensity upon titration with Frag D, and residue-specific response showed that the MP, a continuous stretch of 29 amino acids in PR-R7 binds Frag D. Residue-specific analysis of the binding interaction between PR-R7 and Frag D required initial backbone assignment of this peptide. Very limited dispersion of the NMR signals in the proton dimension raised the suspicion that PR-R7 had limited secondary structure in its apo (unbound) state, which subsequently was shown by the chemical shift prediction using Talos+. This limited dispersion led to substantial signal overlap, and full backbone assignment was ultimately achievable with the aid of novel ^13^C-direct detect measurements geared towards such peptides. A very careful, manual analysis of the titration experiments ensured true representation of the peak intensities for each residue.

Based on the titration results from PR-R7 with Frag D (**Fig. 9**) it is clear that the primary interaction of Frag D is with the residues 20 to 49 on PR-R7. The C-terminal end of the interaction site is defined more clearly than the N-terminal end, although alanine scanning identified both to be required for binding. Both ends are flanked by prolines, and at this point their role at either side of the MP is unclear. Residues 51 to 75 were significantly less impacted by the presence of Frag D, but a secondary interaction with those residues cannot be ruled out as indicated by the alanine scan of that portion of the peptide. The HADG sequence, and aligned YNDG/HANG/HGNG in other repeats may actually participate in subsequent R/R binding of Frag D/Fbg.

A previous study used SC from *S. aureus* strain 8325-4 (22), with a PR and five repeats, for investigation of C-terminal Fbg interactions (28). Enzyme-linked immunosorbent assays and microcalorimetry respectively demonstrated Fbg and Frag D binding to recombinant fragments of the C-terminal SC repeats. Sequence similarities in the SC PR and R1-R2 junction with another *S. aureus* virulence factor, the extracellular fibrinogen-binding protein (Efb) suggested a common Fbg binding motif, absent in the virulence factor von Willebrand factor binding protein (vWbp) that uses another mechanism to recruit Fbg (29). Efb uses Fbg binding to facilitate phagocytosis escape (13), and the pathogen co-opts host Fbg binding by both SC and Efb to evade the host immune system. In ELISA competition assays, short peptides inhibited binding of a C-terminal SC fragment to Fbg-coated microtiter wells. This fragment included the PR and repeats R1 to R5. The competitive peptides were Coa-R0 which has a 32-residue sequence bridging PR and R1; the 27-residue peptides Coa-RI, -RI2, -RI3 and -RI4, all bridging R1 and R2; and Coa-RV1 with a partial R5 sequence. The sequence of repeat R5 in SC 8325-4 largely corresponds to that of repeat R7 in SC Tager 104. Binding of Coa-RV1 to immobilized Fbg and to Frag D by isothermal titration calorimetry cannot readily be explained by potentially different exposure of Frag D epitopes in conformationally heterogeneous immobilized Fbg (51) and in solution. However, three other peptides, Coa-RV2, -RV3 and -RV4 largely corresponding to R5 were not competitive, consistent with our hypothesis that most single repeats may not have a high affinity for Frag D. Binding of Coa-R1 and -RI3 to Frag D, measured by isothermal titration calorimetry yielded affinities between ~77 and ~140 nM, in agreement with our fluorescence equilibrium binding data for the affinities of labeled PR-R constructs (**Table 1**). Ala scanning of ET**<u>N</u>** A**<u>Y</u>** N**<u>V</u>TTH** A**NG** QVS**<u>Y</u>G** A**<u>R</u>P** TYKKPS bridging R1 and R2 in Coa-RI (28) indicated critical (*bold, underlined*) and important (*bold*) residues. Our Ala scanning encompassed a different inter-domain sequence, identified as the MP, comprised of the C-terminal VKYRDA<u>G</u>TG<u>I</u>REYND<u>G</u>TFG<u>Y</u>E residues of the PR, and the N-terminal part **<u>ARP</u>TYK<u>KP</u>** SETN<u>A</u>YN<u>V</u>TT**HAD<u>G</u>**TAT<u>Y</u>G of repeat R7. The alignment of conserved residues in the PR and the repeat junctions (*underlined*) is shown in **Fig. 13**. Conserved residues in the PR proved essential for Frag D binding, and R7 residues in bold were also required (**Fig. 11**). HADG is conserved in the last repeat of the serotypes (49), and aligns with HANG in Coa-RI. Ala scanning of the PR sequence EYNDG^37^ completely abolished Fragment D binding to MP (**Fig. 11**), suggesting that alignment of G^37^ with the conserved G residues in the inter-repeat sequences plays a critical role. PR residues G/A^28^, I/V^31^, G^37^ and Y^41^ are aligned with the C-terminal sequences in the individual repeats. Ko *et al*. (28) identified essential N, Y, T, H and G residues in SC 8325-4 repeat R1, flanking the aligned PR residues, and these may represent a complementary, conserved binding motif in the repeat junctions.

Although competitive ELISA binding to immobilized full-length Fbg may detect additional secondary binding sites for the SC C-terminal repeat region, it is clear that Frag D binding represents a major site of interaction on Fbg. Multiple Frag D binding to SC may indicate the potential of multiple full-length Fbg binding and a role in establishing conformational interactions that may favor Fbg binding in the substrate mode to the complex of SC and host ProT.

Semiquantitative native PAGE analysis suggested stoichiometries for Frag D binding largely consistent with the number of inter-repeat sequences containing conserved motifs, i.e. ~3 for the labeled PR-R1R6R7 and PR-R1R2R3 constructs, and ~7 for labeled PR-(R1→R7) (**Figs. S3, 5**). Quantitative fluorescence equilibrium binding titrations confirmed stoichiometries of 2.5 - 3.3 for all constructs with 3 inter-repeats, and gave stoichiometries of 4.8 - 5.2 for three independent PR-(R1→R7) preparations (**Fig. 15, Table 2**). The *K*_D_ values for Frag D binding to the repeat constructs represent affinities based on the assumption of independent, non-cooperative binding. *K*_D_ values ranged from ~7 to ~32 nM for constructs containing the naturally occurring PR-R1 sequence, and were only marginally higher (~42 nM) for those containing a PR-R3 sequence. Similarly, modest differences were observed for Frag D binding to labeled PR-R1 and PR-R3 constructs containing only one repeat (~50 and ~100 nM, respectively, **Table 1**). Multiple PR-R1R2R3 and [5F]PR-(R1-R7) preparations exhibited acceptable variability in affinities, with good consistency in the stoichiometry and amplitude parameters. This variability is to be expected in nonlinear regression analysis with no constraint imposed on any of the parameters, and using multiple independent protein preparations ([5F]PR-(R1→R7), PR-R1R2R3 and Frag D) in direct and competitive titrations. Minor differences in sample preparation, and limitations of assay sensitivity and of fitting interactions with high stoichiometry factors may all contribute to this variability. Frag D affinities for constructs with multiple repeats were higher than those for PR-R constructs containing only one inter-repeat sequence, suggesting a cumulative effect. The naturally occurring PR-R1 sequence appears to correlate with modestly tighter binding, and the slight sequence variability of the individual repeats might affect the affinity somewhat. However, binding of Frag D to our constructs irrespective of the order of repeats is consistent with the recurring pattern of conserved residues in the inter-repeat regions (**Fig S1**). All PR-three repeat constructs consistently showed a Frag D binding stoichiometry of 1:3, both by native PAGE and fluorescence equilibrium binding, whereas the PR-(R1→R7) constructs appeared to bind ~7 Frag D molecules by native PAGE and ~5 by equilibrium binding. This small discrepancy may reflect limitations of fitting by an unconstrained equilibrium binding model with high ligand stoichiometry, and different conformational binding site constraints in the higher order Frag D-repeat complexes under both experimental conditions.

The Frag D subunit is considerably smaller than Fbg/Fbn, and *in vivo* maximizing Fbg/Fbn occupancy of the SC inter-repeat sequences with minimal sterical hindrance may require a highly organized molecular arrangement such as found in fibrin networks. These networks serve as anchoring points and concealing structures for *S. aureus* to evade the host immune system (28,52). Capturing circulating host Fbg by the C-terminal repeat domain may provide a mechanism of recruiting Fbg for presentation as a substrate of the SC(1-660)·ProT* complex *in vivo*, however Fbg binding to the complex as a substrate for activated ProT* may occur independently of binding to the C-terminal domain of SC. In turbidity assays of Fbg cleavage (**Fig. 14**), Fbg cleavage by thrombin (control) and by the SC(1-325)·ProT* complex was rapid, compared to Fbn formation by the SC(1-660)·ProT* complex at a similar Fbg substrate concentration (**Fig. 14, *d***). Binding to the C-terminal SC domain may sequester the available Fbg for cleavage by the SC(1-660)·ProT* complex, resulting in decreased rates compared to those observed for the SC(1-325)·ProT* complex, lacking the C-terminal domain. Rates and amplitudes of Fbn formation by the SC(1-660)·ProT* complex were dependent on the substrate Fbg concentration, consistent with this hypothesis.

Binding multiple ligand molecules is a characteristic property of intrinsically disordered proteins (53) and intrinsically disordered protein regions (IDPRs) (54). Our CD analysis showed an extended or random secondary structure confirmation (**Fig. 16**) of PR-R7 and PR-(R1→R7) constructs, and secondary structure prediction of full-length SC identified major IDPRs in the C-terminal repeat region (39–41). The presence of disordered structure facilitates binding of multiple ligands, consistent with our observations of 3 and 5 molecules of Frag D binding to PR-three repeat and PR-(R1→R7) constructs, respectively. Our data for binding of multiple Frag D molecules to the repeat constructs did not suggest positive or negative cooperativity, and were fit very well by simple quadratic or cubic binding equations. Although secondary interactions with other Fbg/Fbn domains remain possible, Frag D-mediated binding of multiple Fbg or Fbn molecules in staggered arrays to the disordered SC repeats may induce transition to complexes with a more ordered structure that favor formation of fibrin/bacteria vegetations *in vivo*.

In conclusion, we demonstrated a clear correlation between the specific, highly conserved inter-repeat sequences in the C-terminal domain of SC and the stoichiometry of Frag D binding to these repeat sequences. Stoichiometry assessments determined by fluorescence anisotropy of direct and competitive binding of Frag D to the repeat constructs were in good agreement with those determined by native PAGE. The disordered nature of the C-terminal SC domains may favor identification of conserved linear epitopes for developing therapeutic antibodies to combat *S. aureus* associated infections. These antibodies may be effective across the complete spectrum of serotypes.

## Experimental procedures

### Preparation of Frag D from human Fbg

Frag D was purified from human Fbg as described (55), with some modifications. Fbg (400 mg) was dissolved in 150 mM NaCl, 50 mM imidazole buffer, pH 7.2, to a final concentration of 5-10 mg/ml and dialyzed against the same buffer. CaCl_2_ was added to a final concentration of 5 mM. The Fbg solution was treated with 5 mM iodoacetamide for 15 minutes at room temperature to inhibit cross-linking by trace FXIIIa. Fbg was digested by trypsin (0.01 mg trypsin/mg Fbg) for 4 hours at room temperature. The reaction was terminated by adding soybean trypsin inhibitor (SBTI) to a concentration of 0.03 mg/mg Fbg and incubating for 1 hour on ice. The digested solution was precipitated by slowly adding ice-cold ammonium sulfate solution to a final concentration of 1.18 M, and stirring for 10 minutes on ice. The solution was centrifuged at 10,600 × *g* for 1 hour at 4 °C and the pellet was discarded. The supernatant was further precipitated by slowly adding ice-cold ammonium sulfate solution to a final concentration of 1.61 M and stirring for 10 minutes. The solution was centrifuged for 10,600 × *g* for 1 hour. The pellet was resuspended in 20 mM TRIS buffer, pH 8.0, containing 10 μM *D*-Phe-Phe-Arg-chloromethyl ketone (FFR-CK), 10 μM *D*-Phe-Pro-Arg-chloromethyl ketone (FPR-CK) and 100 μM 4-benzenesulfonyl fluoride hydrochloride (AEBSF), and dialyzed against the same buffer at 4 °C. The dialyzed solution was loaded onto a pre-equilibrated 1 mL Resource Q column, washed with 10 column volumes of 20 mM TRIS buffer, pH 8.0, and eluted with a 20 mM TRIS, 0 - 250 mM NaCl gradient, pH 8.0. Fractions containing Frag D were pooled and concentrated. The solution was dialyzed against 20 mM sodium phosphate, 150 mM NaCl, pH 7.0 (for NMR); or 50 mM HEPES and 125 mM NaCl buffer, pH 7.4 (all other experiments) and stored at −80 °C until use. The Frag D concentration was calculated using M_r_ 88,000 and A_280_ extinction coefficient of 2.08 mL mg^−1^ cm^−1^ (56).

### Preparation of SC C-terminal repeat variants

Combination constructs of PR with single and multiple repeats were amplified by PCR from the SC(1-660) gene of *S. aureus* Newman D2 Tager 104 using degenerate and specific primers, and cloned into a modified pET 30b(+) (Novagen) expression vector containing an N-terminal His_6_-tag followed by a Tobacco Etch Virus (TEV) cleavage site (15,57), and the leading sequence SETTEASHYP. PCR was performed in 50 μL reaction mixtures containing ~10 ng of SC(1-660) template DNA, 1 ng of primer, 0.8 mM dNTPs, and 1 unit of high-fidelity polymerase (Stratagene). A C-terminal Cys residue for 5-IAF labeling was introduced by site-directed mutagenesis (QuikChange) in constructs for fluorescence binding, and all sequences were confirmed by DNA sequencing. The constructs were transformed into *coli* Rosetta 2 (DE3) pLysS. Positive colonies were identified by DNA sequencing. Luria broth (LB, 150 mL containing 0.1 mg/mL of kanamycin) was inoculated with single colonies, and cultured overnight. LB media containing 0.1 mg/mL of kanamycin were inoculated (1:40) with the overnight cultures and grown for 2-3 hours with shaking at 250 RPM and 37 °C until A_600_ of 0.6 was reached. After induction with D-lactose (10 mg/ml) and an additional 4 hour growth, cells were centrifuged at 5000 × *g* for 30 min and lysed in 50 mM HEPES, 125 mM NaCl, 1 mg/mL PEG, 1 mM EDTA buffer, pH 7.4, containing 100 μM phenyl methyl sulfonyl fluoride (PMSF), and 10 μM FFR- and FPR-CK, by three freeze-thaw cycles in liquid N_2_. The lysate was centrifuged at 39,200 × *g* for 45 minutes, and the constructs were released from the inclusion bodies in the pellet by suspension in the same buffer containing 3 M NaSCN. The solution was centrifuged, and dialyzed against 50 mM HEPES, 325 mM NaCl, 50 mM imidazole buffer, pH 7.4 before chromatography on Ni^2+^-iminodiacetic acid-Sepharose (5 ml), and elution with a gradient of 0 - 500 mm imidazole (19). The N-terminal His_6_-tag was removed by incubation with recombinant TEV protease (15,58,59). The C-terminal Cys thiol groups of unlabeled peptides for competitive binding were blocked with 5-iodoacetamide. Peptides (2-4 mg) were initially reduced by adding 2 mM DTT for 30 minutes on ice, and further treated with 4-fold molar excess iodoacetamide for 2 hours in the dark at 20 °C. Excess blocking agent was removed by dialysis against 50 mM HEPES, 125 mM NaCl, pH 7.4. Because the experimentally determined molar extinction coefficients *ε*_280_ of the constructs differed by ~20 % from the theoretical values, molar absorptivities *ε*_205_ of the peptides at 205 nm were determined directly from their amino acid sequence (https://spin.niddk.nih.gov/clore/), and by using measured A_205_ and A_280_, good estimates for *ε*_280_ = *ε*_205_(A_280_/A_205_) were obtained (60). The peptide concentrations were determined from the A_280 nm_ absorbance using molar extinction coefficients *ε*_280_ of 10,430 M^−1^ cm^−1^ for PR-R1, PR-R2, PR-R3; 11,920 M^−1^ cm^−1^ for PR-R6 and PR-R7; 17,880 M^−1^cm^−1^ for PR-R1R2R3 and PR-R1R6R7; 19,370 M^−1^cm^−1^ for PR-R3R4R7 and PR-R3R6R7; and 31,290 M^−1^cm^−1^ for PR-(R1→R7).

Purified peptides destined for labeling were reduced with dithiothreitol (DTT) at 1 mM final concentration, and dialyzed against 5 mM MES, 150 mM NaCl buffer, pH 6.0, before labeling with 5-iodoacetamidofluorescein (5-IAF). Cys-thiol incorporation was measured as described (15). The pH of peptide solutions was raised to ~7 by addition of 0.1 volume 1 M HEPES, pH 7.0, in the presence of a 2-fold molar excess of 5-IAF. After addition of 0.9 volumes of 100 mM HEPES, 100 mM NaCl buffer, pH 7.0, and incubation for 1-2 hours at 25°C, the reaction was quenched by adding DTT to a final concentration of 1 mM. and the mixture was centrifuged for 15 min at 20,820 × *g*. Excess probe was removed by dialysis against 5 mM MES, 150 mM NaCl buffer, pH 6.0. The labeled peptides were further purified by C-18 (5 μm, 4.6 mm × 150 mm, Beckman Coulter Inc.) reverse phase-high performance liquid chromatography. Peptides were eluted using a linear gradient of 0.1 % trifluoroacetic acid (TFA) and 100 % acetonitrile with 0.1 % TFA. The absorbance was measured at 205 nm (peptide bond) and 440 nm (5-fluorescein [5F] probe absorbance). Peak fractions corresponding to 205 nm and 440 nm were pooled and lyophilized. The lyophilized powder was resuspended in water and the labeling ratio was measured. Incorporation of fluorescein was determined by absorbance at 280 nm and 498 nm with an absorption coefficient of 84,000 m^−1^ cm^−1^ for fluorescein and an *A*_280 nm_/*A*_498 nm_ ratio of 0.19 in 6 m guanidine, 100 mm Tris-Cl, 1 mm EDTA buffer, pH 8.5 (61) and determined to be 0.75 to 1.1 mol fluorescein/mol of peptide. [5F]PR, [5F]R1, and [5F]MP were synthesized by Anaspec Inc. The concentrations of [5F]PR, [5F]R1, and [5F]MP were calculated using the molar extinction coefficients *ε*_280_ 5,960; 2,980; and 5,120 M^−1^ cm^−1^, respectively, as determined above (60).

The full-length staphylocoagulase gene SC(1-660) containing a His_6_-tag and TEV cleavage site at the N-terminus was amplified using specific primers from genomic DNA of *S. aureus* Newman D2 Tager 104 and cloned into the same pET30b(+) vector. His_6_-SC(1-660) was expressed in *E. coli* Rosetta 2 (DE3) pLysS, and purified on Ni^2+^-iminodiacetic acid-Sepharose, followed by removal of the His_6_ tag by TEV protease cleavage. The concentration of SC(1-660) was measured using an experimentally determined extinction coefficient of 0.989 M^−1^ cm^−1^ and a molecular mass of 74,390 Da. Our previously prepared SC(1-325) construct was also used in the current study (19,30).

### Expression and purification of ^15^N PR-R7 peptides

Single colonies of Rosetta 2 (DE3) pLysS containing the ^15^N PR-R7 construct were inoculated into 100 mL sterile LB media containing 0.1 mg/mL kanamycin, and grown overnight at 37 °C with shaking at 250 RPM. Overnight cultures (60 mL) were centrifuged for 15 minutes and the pellets were resuspended in 50 mL sterile milliQ water. Sterile flasks containing 900 mL of milliQ water (2 per construct) were each inoculated with 25 mL of resuspended culture. BioExpress cell growth media (Cambridge Isotope Laboratories Inc., Andover, MA) was added (2×100 mL of 10X U-15N, 98 % for the ^15^N peptide), and cultures were grown at 37 °C with shaking at 250 RPM for 2-3 hours in presence of 0.1 mg/mL of kanamycin until an absorbance reading of 0.6 at 600 nm was reached. Peptide expression was induced by adding 10 mg/mL D-lactose and the cultures were grown for 4 more hours at 37 °C with shaking at 250 RPM. Cultures were centrifuged and the peptides were purified by HPLC as described above.

### Expression of PR-R7 alanine constructs

Wild-type PR-R7 with a N-terminal His_6_ tag followed by a TEV cleavage site, and an engineered C-terminal Cys residue, cloned into the pET30b(+) vector, was used as a starting construct. Residues 22 - 49, constituting the MP region, and the subsequent R7 residues 50 - 69 were converted group-wise (4 to 5 residues, consecutively) to alanine residues through QuikChange site-directed mutagenesis and confirmed by DNA sequencing. The constructs were transformed into *E. coli* Rosetta 2 (DE3), expressed, purified by His_6_ tag and TEV cleavage, labeled with 5-fluorescein, and purified by HPLC as described above. The peptide concentration was determined from the A_280 nm_ absorbance using molar extinction coefficients of 10,430 M^−1^ cm^−1^ for PR-R7 constructs with sequences VKYRDA, EYNDG, TFGYE, YKKP, YNVTT, TATYG changed to alanine residues; and 11,920 M^−1^ cm^−1^ for PR-R7 constructs with sequences GTGIR, ARPT, SETNA, HADG changed to alanines.

### Size exclusion chromatography and light scattering

Size exclusion chromatography demonstrated that full-length SC(1-660), but not SC(1-325), interacts with Frag D. The size exclusion profile was recorded using a Superdex-200 column (GE, 10 × 300 mm) pre-equilibrated with 50 mM HEPES, 125 mM NaCl, 5mM CaCl_2_, pH 7.4 buffer. In a first set of experiments, 100 μL each of the samples of SC(1-325) (52.67 μM or 2.0 mg/mL); Frag D (24.7 μM or 2.1 mg/mL); and 100 μL of a 1:1 mixture of SC(1-325) and Frag D in 50 mM HEPES and 125 mM NaCl buffer, pH 7.4, were passed on the column separately. Subsequently, 100 μL each of the samples of SC(1-660) (24.2 μM or 1.8 mg/mL); Frag D (24.7 μM / 2.1 mg/mL); and 100 μL of a 1:1 mixture of SC(1-660) and Frag D in 50 mM HEPES and 125 mM NaCl, pH 7.4, were passed separately on the pre-equilibrated column and eluted using the same buffer as described above. Elution profiles (absorbance A_280 nm_ in mOD *vs.* elution volume) were recorded continuously for both sets. Light scattering data were acquired with a PD2010 multi-detection light scattering system to obtain the apparent molecular mass (M_r_) of the eluted complexes and proteins.

### Native PAGE of Frag D binding to SC(1-660), [5F]PR-R1, the SC(1-660)·[5F]ProT complex, and labeled C-terminal repeat constructs

SC(1-660) (4 μM) was incubated with 1 to 9x molar excess of Frag D for 30 minutes in 50 mM HEPES, 110 mM NaCl, 5 mM CaCl_2_, 1 mg/mL PEG-8000 containing 10 μM FFR-CK and FPR-CK at 25 °C, and run on native PAGE along with Frag D and SC(1-660) as controls (**Fig. 3C**). The samples were mixed with native sample buffer, 0.32 M Tris, 50 % glycerol and 0.1 % bromophenol blue, pH 7.6, and electrophoresed at 100 volt for 5 - 6 hours at 4 °C on a 6 % Tris-glycine gel without sodium dodecyl sulfate (SDS), using a 0.025 mM Tris, 0.192 mM glycine running buffer, pH 8.3. Gels were stained for protein with GelCode Blue (Pierce), and fluorescence was visualized with a 300 nm transilluminator.

Mixtures of [5F]PR-R1 (12.2 μM), [5F]R1 (50.0 μM) or [5F]PR (45.0 μM) with ~0.5- to ~2.0-fold molar excess of Frag D were incubated and electrophoresed similarly (**Fig. S2**).

[5F]ProT (2.5 μM) was incubated with SC(1-660) (3.7 μM) for 30 minutes, and the complex was reacted with Frag D at 7.5, 11.2, 15.0, 18.7, 26.2 and 29.9 μM (**Fig.4**). [5F]PR-(R1→R7) (4.3 μM) was incubated with Frag D at 10.0, 15.0, 20.0, 25.0, 30.0, 35.0 and 40.0 μM for 30 minutes and electrophoresed similarly (**Fig. 5**).

[5F]PR-R1R6R7 (5.6 μM) and [5F]PR-R1R3R3 (8.1 μM) were incubated with Frag D at 5.0, 10.0, 20.0, 30.0, 40.0, 50.0 and 60.0 μM, and electrophoresed similarly (**Fig. S3**).

### NMR spectroscopy, data collection, and chemical shift assignments

Titration NMR experiments were performed at 25 °C on a Bruker AV-III 600.13 MHz spectrometer equipped with quadruple resonance cryogenically cooled CPQCI probe. Standard ^1^H-^15^N heteronuclear single quantum coherence (HSQC) experiments were used, containing a flip-back pulse and Watergate sequence to suppress the water signal (62–64). Titration experiments were done using 200 μL of 65 μM protein solution plus 5 μL D_2_O in a 3 mm NMR tube. Various amounts of a 240 μM Frag D stock in the same buffer solution were added. All experiments were referenced against the 4,4-dimethyl-4-silapentane-1-sulfonic acid (DSS) standard, processed with Topspin 3.5 (Bruker BioSpin, Billerica, MA) and analyzed with NMRView (One Moon Scientific, Inc., Westfield, NJ). Backbone resonance assignments were utilized as described in (27).

### Fluorescence equilibrium binding of C-terminal repeat peptides to Frag D

Fluorescence intensity and anisotropy measurements were performed with a QuantaMaster 30 spectrofluorometer (Photon Technology International). Titrations were performed at 25 °C in 50 mM HEPES, 110 mM NaCl, 5 mM CaCl_2_ and 1 mg/mL PEG-8000, pH 7.4 buffer with 10 μM FPR-CK inhibitor, at λ_ex_ 490 nm (1-2 nm band pass) and λ_em_ 514 nm (1-5 nm band pass) using acrylic cuvettes coated with PEG 20,000. Fluorescein-labeled SC C-terminal repeat constructs, [5F]PR, [5F]R1, [5F]MP, [5F]PR-R1, [5F]PR-R2, [5F]PR-R3, [5F]PR-R6, [5F]PR-R7, and the Ala-mutated constructs [5F]PR-R7-VKYRDA, [5F]PR-R7-GTGIR, [5F]PR-R7-EYNDG, [5F]PR-R7-TFGYE, [5F]PR-R7-ARPT, [5F]PR-R7-YKKP, [5F]PR-R7-SETNA, [5F]PR-R7-YNVTT, [5F]PR-R7-HADG and [5F]PR-R7-TATYG were titrated with Frag D. The fractional change in fluorescence was calculated as (*F*_obs_ − *F*_o_)/*F*_o_ = *ΔF*/*F*_o_, and the data were fit by the quadratic binding equation (Eq. 2) (65). Nonlinear least-squares fitting was performed with SCIENTIST (MicroMath) to obtain the dissociation constant, *K*_D_, maximum fluorescence intensity ((*F*_max_ − *F*_o_)/*F*_o_ = *ΔF*_max_/*F*_o_), and stoichiometric factor (*n*) for the peptide constructs. The error estimates (2SD) represent the 95% confidence interval. Fitting of the Ala constructs was done with the stoichiometric factor (*n*) fixed to 1, appropriate for titrations with probe concentration below the *K*_D_ of the interaction.

To determine whether the repeats bind to the Frag D β and γ holes, [5F]PR-R1 was titrated with Frag D in the presence of 4.9 mM GPRP under experimental conditions described above; and the *K*_D_ and maximum fluorescence intensity were obtained by fitting the data with *n* fixed to 1.

To determine the effect of pH and NMR buffer conditions, equilibrium binding at pH 7.0 and 7.4 of [5F]PR-R7 (17 and 848 nM, respectively) to Frag D was measured in 20 mM sodium phosphate, 150 mM NaCl, and 1 mg/mL PEG-8000 buffers containing 10 μM FPR-CK inhibitor, using λ_ex_ 490 nm (4 nm band pass at pH 7.0; 2 nm at pH 7.2 and 7.4) and λ_em_ 514 nm (8 nm band pass at pH 7.0; 4 nm at pH 7.4). The *K*_D_ and maximum fluorescence intensity values were obtained as described above, with *n* fixed to 1.

Fluorescence anisotropy titrations were performed in the same buffer. Anisotropy measurements were corrected for total intensity changes using a modified Lakowicz equation (66). [5F]PR-(R1→R7) was titrated with Frag D at excitation wavelength λ_ex_ 495 nm and emission wavelength λ_em_ 514 nm with band passes ranging from 2-6 nm depending on the [5F]PR-(R1→R7) concentration. Three different [5F]PR-(R1→R7) preparations were titrated with three different Frag D preparations to show reproducibility (**Fig. 15, A1, B1, C1**). Measurements of *r*_obs_ as a function of Frag D concentration were analyzed by the quadratic binding equation (65) to obtain the dissociation constant *K*_D_ and stoichiometric factor *n* of ligand binding, initial *(r*_o_) and maximal *(r*_max_) anisotropies, and the change in anisotropy (Δ*r* = *r*_max_ -*r*_o_).

In competitive titrations, mixtures of a fixed concentration of [5F]PR-(R1-R7) with two different concentrations of Frag D were each titrated with unlabeled PR-R1R2R3 (3 independent batches), PR-R1R6R7, PR-R3R4R7 and PR-R3R6R7 to obtain two data curves for each competitor (**Fig. 15, A2, B2, B3, C2, C3, C4**). Concentrations of [5F]PR-(R1-R7) and Frag D in the competitive titrations are given in the figure legends. Respective sets of direct and competitive titrations were analyzed simultaneously by nonlinear least-squares fitting of the cubic equation in SCIENTIST (MicroMath) (65,66) to obtain the *K*_D_ and stoichiometric factor *n*, initial *(r*_o_) and maximal *(r*_max_) anisotropies, and the maximal change in anisotropy (Δ*r* = *r*_max_ - *r*_o_) for competitor binding to Frag D. Error estimates (±2SD) represent the 95% confidence interval.

### Clotting of Fbg by the SC(1-325)·ProT^QQQ*^ and SC(1-660)·ProT^QQQ*^ complexes

Fbg cleavage by thrombin, and the SC(1-325)·ProT^QQQ^* and SC(1-660)·ProT^QQQ^* complexes was measured as described previously (59). Assays were performed in 50 mM HEPES, 110 mM NaCl, 5 mM CaCl_2_, 1mg/ml PEG-8000, pH 7.4 reaction buffer. Increases in turbidity were measured at 450 nm, 25 °C in a microtiter plate reader, for mixtures of Fbg (0.5 mg/mL) with 10 nM thrombin or 10 nM SC(1-325)·ProT^QQQ*^ complex; and mixtures of Fbg at increasing concentrations (0.3, 0.5, 0.75 and 1.0 mg/mL) with 10 nM SC(1-660)·ProT^QQQ*^ complex. Reactions were started by adding thrombin to the Fbg/buffer mixture; or by adding ProT^QQQ^ to the mixtures of Fbg, SC (10 nM) and buffer. Due to the very tight binding of SC(1-325) and SC(1-660) to ProT^QQQ^, the complex concentrations were estimated to be ~10 nM. Control reactions were performed without thrombin or ProT^QQQ^.

### Circular dichroism spectroscopy of SC repeat peptides and Frag D

CD spectra of PR-R7 (10.09 μM), PR-(R1→R7) (3.3 μM), SC(1-325) (2.63 μM), SC(1-660) (1.37 μM), and Frag D (3.93 μM) were measured using a Jasco J-810 CD spectrometer from Beckman Coulter Inc. The spectra were measured in 20 mM sodium phosphate and 150 mM NaCl, pH 7.0 buffer, at peptide/protein concentrations of 0.1 mg/ml. Solutions were inspected for absence of turbidity. Spectra were measured from 260 to 200 nm with high sensitivity, band width set to 1 nm, response time 2 sec and scanning speed of 50 nm/min at 25 °C. Each scan represents the average of three individual scans. Data between 190 and 200 nm were noisy, prohibiting accurate analysis in this region. Analysis was performed as described (67) using the formula, [θ] = (100(signal))/cnl, where [θ] is the mean residue ellipticity in deg cm^2^ decimol^−1^, signal is raw output in millidegrees, c is the peptide or protein concentration in mM, n is the number of amino acids, and l is the cell path length in cm. Ellipticity data were plotted without using a smoothing algorithm. GOR IV, PSIPRED and PONDR-FIT secondary structure predictions were performed online as described (39–41).

## Data availability

The NMR data set “PR-R7 from staphylocoagulase of *S. aureus* Newman D2 Tager 104 strain” has been deposited in the Biological Magnetic Resonance Data Bank, with entry ID 27036: https://bmrb.io/data_library/summary/index.php?bmrbId=27036 All other data are contained within the article and Supporting Information.

## Funding and additional information

This research was supported in part by NIH R01 HL071544 to PEB and IMV, R01 HL114477 to PP; and R01 GM080403 to JM. The NMR instrumentation used in this work was supported by NIH S10 RR026677 and NSF DBI-0922862. The content is solely the responsibility of the authors and does not necessarily represent the official views of the National Institutes of Health.

## Footnotes

### Abbreviations are

*S. aureus*: *Staphylococcus aureus*
ProT: prothrombin
T: thrombin
SC: staphylocoagulase
SC(1-660): full-length SC
Fbg: fibrinogen
Fbn: fibrin
Frag D: fibrinogen fragment D
R1 to R7: repeat 1 to 7
PR: pseudo-repeat
GPRP: Gly-Pro-Arg-Pro
[5F]: [5-fluorescein]

